# A direct and widespread role for the nuclear receptor EcR in mediating the response to ecdysone in *Drosophila*

**DOI:** 10.1101/517458

**Authors:** Christopher M. Uyehara, Daniel J. McKay

**Affiliations:** Department of Biology, Chapel Hill, NC, 27599; Department of Genetics, Chapel Hill, NC, 27599; Curriculum in Genetics and Molecular Biology, Chapel Hill, NC, 27599; Integrative Program for Biological and Genome Sciences, Chapel Hill, NC, 27599; The University of North Carolina at Chapel Hill, Chapel Hill, NC, 27599

**Author notes:** Corresponding author: Daniel J. McKay, Assistant Professor, Department of Biology, Department of Genetics, Integrative Program for Biological and Genome Sciences, The University of North Carolina at Chapel Hill, Chapel Hill, NC 27599, USA.

**Keywords:** Hormone, Transcription Factor, CUT&RUN

## Abstract

**ABSTRACT:** The ecdysone pathway was amongst the first experimental systems employed to study the impact of steroid hormones on the genome. In *Drosophila* and other insects, ecdysone coordinates developmental transitions, including wholesale transformation of the larva into the adult during metamorphosis. Like other hormones, ecdysone controls gene expression through a nuclear receptor, which functions as a ligand-dependent transcription factor. Although it is clear that ecdysone elicits distinct transcriptional responses within its different target tissues, the role of its receptor, EcR, in regulating target gene expression is incompletely understood. In particular, EcR initiates a cascade of transcription factor expression in response to ecdysone, making it unclear which ecdysone-responsive genes are direct EcR targets. Here, we use the larval-to-prepupal transition of developing wings to examine the role of EcR in gene regulation. Genome-wide DNA binding profiles reveal that EcR exhibits widespread binding across the genome, including at many canonical ecdysone-response genes. However, the majority of its binding sites reside at genes with wing-specific functions. We also find that EcR binding is temporally dynamic, with thousands of binding sites changing over time. RNA-seq reveals that EcR acts as both a temporal gate to block precocious entry to the next developmental stage as well as a temporal trigger to promote the subsequent program. Finally, transgenic reporter analysis indicates that EcR regulates not only temporal changes in target enhancer activity but also spatial patterns. Together, these studies define EcR as a multipurpose, direct regulator of gene expression, greatly expanding its role in coordinating developmental transitions.

**SIGNIFICANCE:** Nuclear receptors (NRs) are sequence-specific DNA binding proteins that act as intracellular receptors for small molecules such as hormones. Prior work has shown that NRs function as ligand-dependent switches that initiate a cascade of gene expression changes. The extent to which NRs function as direct regulators of downstream genes in these hierarchies remains incompletely understood. Here, we study the role of the NR EcR in metamorphosis of the *Drosophila* wing. We find that EcR directly regulates many genes at the top of the hierarchy as well as at downstream genes. Further, we find that EcR binds distinct sets of target genes at different developmental times. This work helps inform how hormones elicit tissue- and temporal-specific responses in target tissues.

## INTRODUCTION

Hormones function as critical regulators of a diverse set of physiological and developmental processes, including reproduction, immune system function, and metabolism. During development, hormones act as long-range signals to coordinate the timing of events between distant tissues. The effects of hormone signaling are mediated by nuclear receptors, which function as transcription factors that differentially regulate gene expression in a hormone-dependent manner. Whereas many of the co-regulators that contribute to nuclear receptor function have been identified, the mechanisms used by these factors to generate distinct, yet appropriate, transcriptional responses in different target tissues are incompletely understood.

Ecdysone signaling has long served as a paradigm to understand how hormones generate spatial and temporal-specific biological responses. In *Drosophila*, ecdysone is produced in the prothoracic gland and is released systemically at stereotypical stages of development (1, 2). Pulses of ecdysone are required for transitions between developmental stages, such as the larval molts. A high titer pulse of ecdysone triggers the end of larval development and the beginning of metamorphosis (1, 2). With each pulse, ecdysone travels through the hemolymph to reach target tissues, where it enters cells and binds its receptor, a heterodimer of the proteins EcR (*Ecdysone receptor,* a homolog of the mammalian Farnesoid X Receptor) and Usp (*ultraspiracle,* homolog of mammalian RXR) (3–5). In the absence of ecdysone, EcR/Usp is nuclear-localized and bound to DNA where it is thought to act as a transcriptional repressor (6, 7). Upon ecdysone binding, EcR/Usp switches to a transcriptional activator (6). Consistent with the dual regulatory capacity of EcR/Usp, a variety of co-activator and co-repressor complexes have been shown to function with this heterodimer to regulate gene expression (7–13).

Understanding how ecdysone exerts its effects on the genome has been heavily influenced by the work of Ashburner and colleagues in the 1970’s. By culturing larval salivary glands *in vitro,* Ashburner described a sequence of visible puffs that appear in the giant polytene chromosomes upon addition of ecdysone (14, 15). A small number of puffs appeared immediately after ecdysone addition, followed by the appearance of more than one hundred additional puffs over the next several hours (14–16). The appearance of early puffs was found to be independent of protein synthesis, suggesting direct action by EcR/Usp, whereas the appearance of late puffs was not, suggesting they require the protein products of early genes for activation (1, 15, 17). These findings, and decades of subsequent work elucidating the molecular and genetic details, have led to a hierarchical model of ecdysone signaling in which EcR/Usp directly induces expression of a small number of early response genes. Many of these early response genes encode transcription factors, such as the zinc finger protein Broad, the nuclear receptor Ftz-f1, and the pipsqueak domain factor E93 (2). The early response transcription factors are required, in turn, to induce expression of the late response genes, which encode proteins that impart the temporal and tissue-specific responses to ecdysone in target tissues.

Although the framework of the ecdysone pathway was established through work in salivary glands, additional studies have affirmed an essential role for ecdysone signaling in many other tissues. Similar to other hormones, the physiological response to ecdysone is often profoundly specific to each target tissue. For example, ecdysone signaling triggers proliferation, changes in cell and tissue morphology, and eventual differentiation of larval tissues that are fated to become part of the adult fly, such as the imaginal discs (2, 18). By contrast, ecdysone signaling initiates the wholesale elimination of obsolete tissues, such as the larval midgut and salivary glands through programmed cell death (2, 18, 19). Ecdysone also has essential functions in the nervous system during metamorphosis by directing remodeling of the larval neurons that persist until adulthood, and in specifying the temporal identity of neural stem cell progeny born during this time (20). While it is clear that ecdysone signaling triggers the gene expression cascades that underlie each of these events, the molecular mechanisms by which ecdysone elicits such diverse transcriptional responses in different target tissues remains poorly understood.

A key step in delineating the mechanisms by which ecdysone signaling regulates target gene expression involves identification of EcR/Usp DNA binding sites. Given the hierarchical structure of the ecdysone pathway, it is unclear if EcR acts primarily at the top of the transcriptional cascade, or if it also acts directly on downstream effector genes. Several early response genes such as *br, Eip74EF,* and the glue genes have been shown to be directly bound by EcR *in vivo* (5, 21, 22). At the genome-wide level, polytene chromosome staining revealed approximately 100 sites bound by EcR in larval salivary glands (23). DamID and ChIP-seq experiments have identified roughly 500 sites directly bound by EcR in *Drosophila* cell lines (24, 25). Thus, the available evidence, albeit limited, indicates that EcR binds to a limited number of target genes, consistent with hierarchical models wherein the response to ecdysone is largely driven by early response genes and other downstream factors.

We recently identified the ecdysone-induced transcription factor E93 as being essential for the proper temporal sequence of enhancer activation during pupal wing development (26). In the absence of E93, early-acting wing enhancers fail to turn off, and late-acting wing enhancers fail to turn on. Moreover, ChIP-seq identified thousands of E93 binding sites across the genome. These data support the hierarchical model of ecdysone signaling in which early response transcription factors like E93 directly regulate a significant fraction of ecdysone-responsive genes in target tissues.

Here, we sought to determine the role that EcR performs in temporal gene regulation during the larval-to-prepupal transition of the wing. Using wing-specific RNAi, we find that EcR is required for proper morphogenesis of prepupal wings, although it is largely dispensable for wing disc patterning at earlier stages of development. RNA-seq profiling reveals that EcR functions as both a temporal gate to prevent the precocious transition to prepupal development as well as a temporal trigger to promote progression to next stage. Using CUT&RUN, we map binding sites for EcR genome wide before and after the larval-to-prepupal transition. Remarkably, we find that EcR binds extensively throughout the genome, including at many genes with wing-specific functions that are not part of the canonical ecdysone signaling cascade. Moreover, EcR binding is highly dynamic, with thousands of binding sites gained and lost over time. Finally, transgenic reporter analyses demonstrate that EcR is required not only for temporal regulation of enhancer activity, but also for spatial regulation of target enhancers. Together, these findings indicate that EcR does not control gene expression solely through induction of a small number of downstream transcription factors, but instead it plays a direct and widespread role in regulating tissue-specific transcriptional programs.

## RESULTS

### Temporal changes in gene expression during the larval-to-prepupal transition

In *Drosophila,* the end of larval development marks the beginning of metamorphosis. Over a five-day period, larval tissues are destroyed, and the progenitors of adult tissues, such as wing imaginal discs, undergo a series of progressive morphological and cell differentiation events to acquire their final shapes and sizes. By the end of larval development, the wing disc is comprised of a largely undifferentiated array of columnar epithelial cells (27, 28). The first 12 hours after puparium formation (APF) is termed the prepupal stage. During this period, cell division is arrested, and the pouch of the wing disc everts outward, causing the dorsal and ventral surfaces of the wing to appose one another, forming the presumptive wing blade (**Fig 1A-B**) (27, 28). At the same time, the notum of the wing disc extends dorso-laterally, and eventually fuses with the contralateral wing disc to form the back of the adult fly (**Fig 1A-B**). Additional events occurring during this time period include secretion of the prepupal cuticle and migration of muscle progenitor cells.

**Figure 1:**
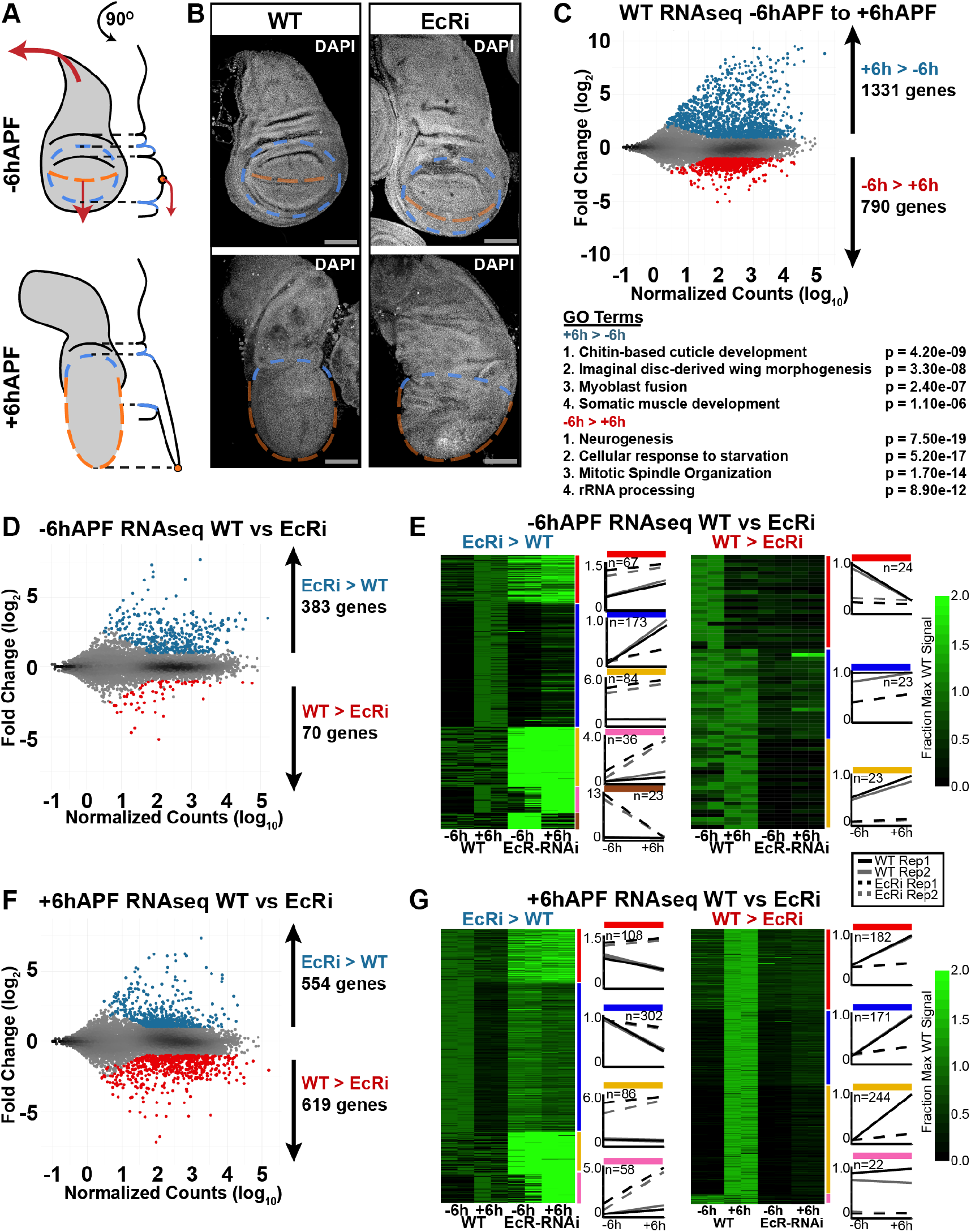
EcR is required to promote global changes in gene expression in wings between - 6hAPF and +6hAPF. (A) Cartoon diagram of wildtype (WT) wing eversion between −6hAPF and +6hAPF. (B) Confocal images of *WT* wings and wings expressing *UAS-EcR RNAi* from *vg-tubGAL4* (hereafter EcR-RNAi) at −6hAPF and +6hAPF. The dorsal-ventral (DV) boundary is marked by an orange dotted line. The edge of the pouch is indicated by a blue dotted line. (B) MA plots (top) and gene ontology terms (bottom) of RNAseq comparing gene WT wings at −6hAPF and +6hAPF. (C-D) MA Plots and clustered heatmaps of RNAseq comparing EcR-RNAi wings and WT wings at −6hAPF. (E-F) MA plots and heatmaps of RNAseq comparing EcR-RNAi wings at WT wings at +6hAPF. Scale bars for immunostaining are 100μm. For MA plots, differentially expressed genes (padj < 0.05, absolute log_2_ fold change > 1) are colored red and blue. Heatmaps are represented as the fraction of max WT counts. Colored bars to the right denote start and end of each cluster. Line plots are the mean signal for each cluster.

To understand EcR’s role in promoting the larval-to-prepupal transition, we began by identifying global changes in gene expression that occur in wild type wings before and after the onset of pupariation. We collected wing tissue from wandering, third instar larvae, approximately six hours prior to puparium formation (hereafter, –6hAPF) and from prepupae, approximately six hours after puparium formation (hereafter, +6hAPF), and performed RNAseq. As described previously (28), wildtype gene expression is highly dynamic during this time period. Using a conservative definition for differential expression (FDR < 0.05, >= 2-fold change in expression), we identified over 1300 genes increasing in expression and nearly 800 genes decreasing in expression (**Fig 1C**). The observed gene expression changes are consistent with developmental events occurring at this time. For example, genes that increase over time are involved in cuticle deposition, wing morphogenesis, and muscle development (**Fig 1C**). By contrast, genes that decrease over time are involved in cell cycle regulation, cellular metabolism, and neural development. Thus, the morphological changes that define the larval-to-prepupal transition are rooted in thousands of changes in gene expression.

### EcR is required for the larval-to-prepupal transition in wings

The onset of pupariation is induced by a high titer ecdysone pulse. At the genetic level, ecdysone acts through its receptor, EcR. Null mutations in *EcR* are embryonic lethal. Therefore, to investigate the role that EcR plays in wing development, we used a wing-specific GAL4 driver in combination with an RNAi construct to knockdown EcR expression throughout wing development (29). EcR-RNAi driven in wing discs diminished protein levels by approximately 95% (**Fig S1A-C**).

In agreement with previous work suggesting that EcR does not appear to be required for wing development during the 1^st^ and 2^nd^ instar stages (30–33), EcR-RNAi wings appear morphologically similar to wild type (WT) wing imaginal discs at –6hAPF (**Fig 1B**). However, EcR-RNAi wing discs are noticeably larger than WT wing discs, consistent with the proposed role for ecdysone signaling in cell cycle inhibition in 3^rd^ instar larvae (30, 31). By contrast, EcR-RNAi wings at +6hAPF appear morphologically dissimilar to both –6hAPF EcR-RNAi wings and to WT wings at +6hAPF. The pouch fails to properly evert and larval folds remain visible. Similarly, the notum fails to extend appropriately, and appears more similar to the larval notum than the notum at +6hAPF (**Fig 1B**). These findings suggest that wings fail to properly progress through the larval-to-prepupal transition in the absence of EcR. Notably, this failure is likely not due to a systemic developmental arrest because legs isolated from larvae and pupae expressing EcR-RNAi in the wing exhibit no morphological defects (**Fig S1D**). We conclude that EcR is required tissue-autonomously for progression through the larval-to-prepupal transition.

To identify genes impacted by the loss of EcR, we performed RNA-seq on EcR-RNAi wings at –6hAPF and +6hAPF. Knockdown of EcR results in widespread changes in gene expression (**Fig 1D**). At –6hAPF, 453 genes are differentially expressed in EcR-RNAi wings relative to wildtype wing imaginal discs. Remarkably, 85% of these genes (n=383, “–6hAPF EcRi > WT”) are expressed at higher levels in EcR-RNAi wings relative to WT, suggesting that EcR is primarily required to repress gene expression at −6hAPF. To determine the expression profiles of these genes during WT development, we performed cluster analysis (**Fig 1E**), and found that 72% of these −6hAPF EcRi UP genes normally increase in expression between −6hAPF and +6hAPF (**Fig 1E**). Genes in this category include those involved in cuticle development as well as multiple canonical ecdysone response genes (**Table S1**). Thus, a major role of EcR at −6hAPF is to keep genes involved in the prepupal program from being precociously activated during larval stages.

We next examined the impact of EcR knockdown in +6hAPF wings. In contrast to the effect at −6hAPF, wherein genes primarily increased in the absence of EcR, we observed approximately equal numbers of up- and down-regulated genes relative to WT wings at +6hAPF (**Fig 1F**). Clustering of EcR-RNAi and WT RNA-seq data revealed distinct differences in the inferred regulatory role of EcR at +6hAPF relative to −6hAPF (**Fig 1G**). 74% of the genes expressed at higher levels in EcR-RNAi wings relative to WT normally decrease in expression between −6hAPF and +6hAPF (**Fig 1G**). Genes in this category include factors that promote sensory organ development and metabolic genes (**Table S2**). The increased levels of these “+6hAPF EcRi > WT” genes suggest that in addition to preventing precocious activation of the prepupal gene expression program, EcR is also required to shut down the larval gene expression program. However, we also observe a role for EcR in gene activation. For genes that are expressed at lower levels in EcR-RNAi wings (n=619, “+6hAPF WT > EcRi”), 96% of these genes normally increase between −6hAPF and +6hAPF. Genes in this category include those involved in muscle development as well as regulators of cell and tissue morphology (**Table S2**). We conclude that EcR is required not only for gene repression but also for gene activation, consistent with the demonstrated interaction of EcR with both activating and repressing gene regulatory complexes (7–13). Collectively, these data demonstrate that the failure of EcR-RNAi wings to progress through the larval-to-prepupal transition coincides with widespread failures in temporal gene expression changes.

### EcR directly binds thousands of sites genome-wide

The experiments described above reveal that ecdysone triggers thousands of gene expression changes in wings during the larval-to-prepupal transition. Because ecdysone signaling initiates a cascade of transcription factor expression, it is unclear which of these changes are mediated directly by EcR. Therefore, we next sought to determine the genome-wide DNA binding profiles of EcR in developing wings. For these experiments, we utilized a fly strain in which the endogenous *EcR* gene product has been epitope-tagged by a transposon inserted into an intron of *EcR* (34, 35). This epitope tag is predicted to be incorporated into all EcR protein isoforms (hereafter EcR^GFSTF^) (**Fig S2A**). Genetic complementation tests determined that *EcR^GFSTF^* flies are viable at the expected frequency (**Fig S2B**), indicating that epitope-tagged EcR proteins are fully functional. Supporting this interpretation, western blotting demonstrated that EcR^GFSTF^ protein levels are equivalent to untagged EcR, and immunofluorescence experiments revealed nuclear localization of EcR^GFSTF^ (**Fig S2C-D**), as well as binding of EcR^GFSTF^ to the same regions as untagged EcR on polytene chromosomes (**Fig S2E**).

To generate genome-wide DNA binding profiles for EcR, we performed CUT&RUN on – 6hAPF wings (**Fig 2A**) from *EcR^GFSTF^* flies. CUT&RUN provides similar genome-wide DNA binding information for transcription factors as ChIP-seq, but requires fewer cells as input material (36, 37), making it useful for experiments with limiting amounts of tissue. Our EcR CUT&RUN data exhibit features that are similar to those previously reported for other transcription factors, including greater DNA-binding site resolution relative to ChIP-seq (**Fig S3, S5**). Wing CUT&RUN profiles at –6hAPF reveal that EcR binds extensively throughout the genome (**Fig 2**). Many EcR binding sites localize to canonical ecdysone target genes, including *broad, Eip93F, Hr3, Hr4 and Eip75B* (**Fig 2A**). Surprisingly, we also observed EcR binding to many genes that have not previously been categorized as ecdysone targets, including *homothorax, Delta, Actin 5C, Stubble and crossveinless c* (**Fig 2B**). Thus, EcR binds widely across the genome in wing imaginal discs. The widespread binding of EcR observed here contrasts with previous genome-wide DNA binding profiles obtained for EcR. For example, ChIP-seq profiles from S2 cells and DamID profiles from Kc167 cells identified 500-1000 binding sites (24, 25). By contrast, our findings demonstrate that EcR binds both canonical and non-canonical ecdysone-target genes, raising the question as to whether EcR directly contributes to a wing-specific transcriptional program.

**Figure 2:**
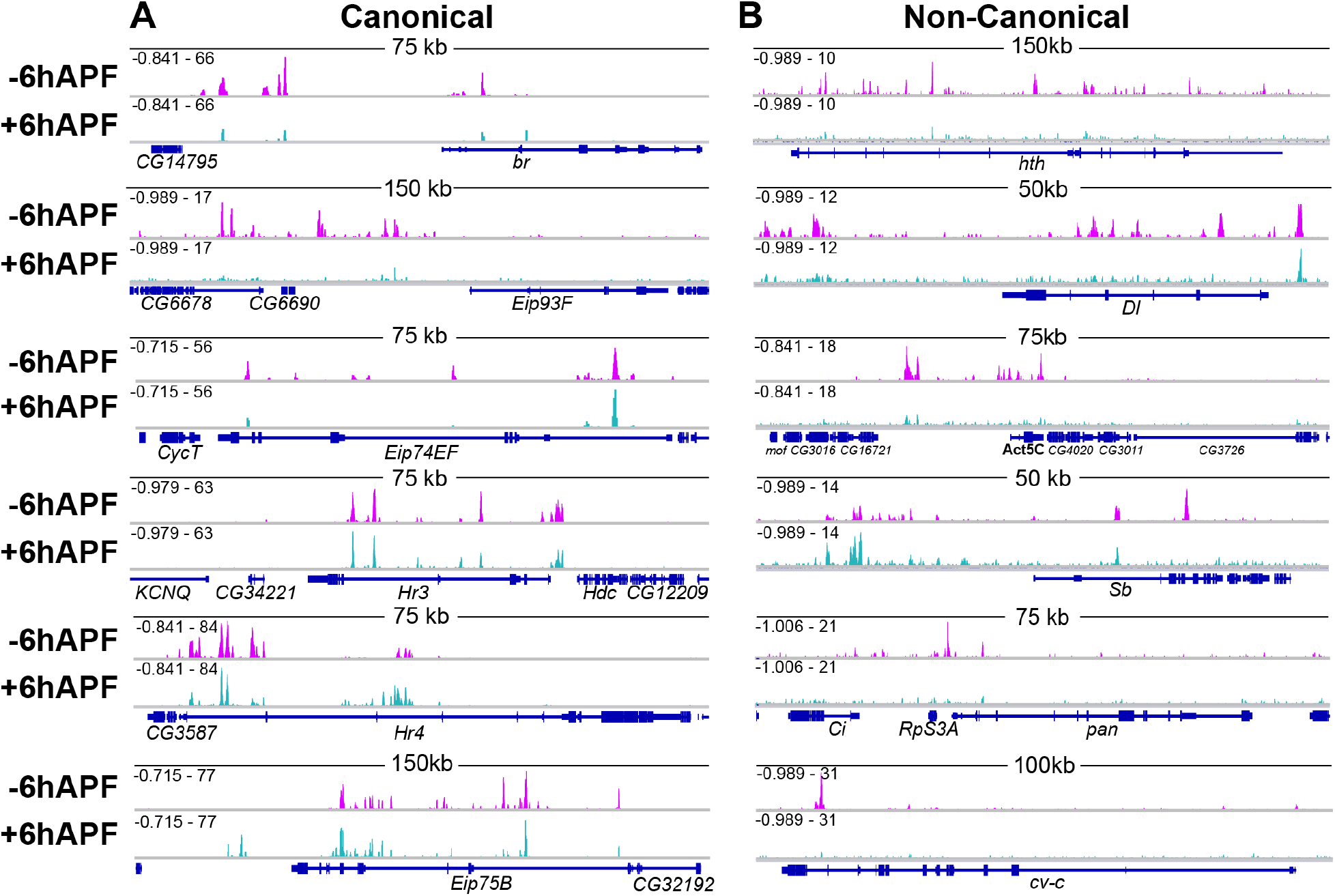
EcR binds extensively throughout the genome. Browser shots of EcR CUT&RUN signal (z-score) at −6hAPF and +6hAPF at (A) canonical and (B) non-canonical ecdysone response genes. Signal range is indicated in top-left corner.

In addition to widespread DNA binding, we also observed clustering of EcR binding sites in the genome. EcR peaks are significantly closer to one another than expected by chance (**Fig S4A-C**), and a majority of peaks are located within 5kb of an adjacent peak (**Fig S4D**). In particular, canonical ecdysone target genes often exhibit clusters of EcR binding (**Fig S4E-F**). These findings suggest that EcR often binds multiple *cis*-regulatory elements across target gene loci, consistent with the observed clustering of ecdysone-responsive enhancers in S2 cells (25).

### EcR binding is temporally dynamic

To understand the role of EcR binding in temporal progression of wing development, we next performed CUT&RUN on +6hAPF wings (**Fig 2, 3A**). Similar to our findings from −6hAPF wings, we found that EcR binds widely across the genome at +6hAPF. Interestingly, there is a global decrease in the number of sites occupied by EcR over time: EcR binds to a total of 4,967 sites genome-wide at −6hAPF, whereas it binds 1,174 sites at +6hAPF (**Fig 3B**). While many of the +6hAPF binding sites overlap with −6hAPF binding sites (763 peaks, 65%) (hereafter −6h/+6h stable binding sites), we also identified 411 sites that are specific to the +6hAPF time. Similar to –6hAPF peaks, EcR binding sites at +6hAPF peaks are clustered genome-wide (**Fig S4**). Thus, the larval-to-prepupal transition in wings is marked by both the loss of EcR from the majority of its – 6hAPF binding sites, as well as the gain of EcR at hundreds of new binding sites.

**Figure 3:**
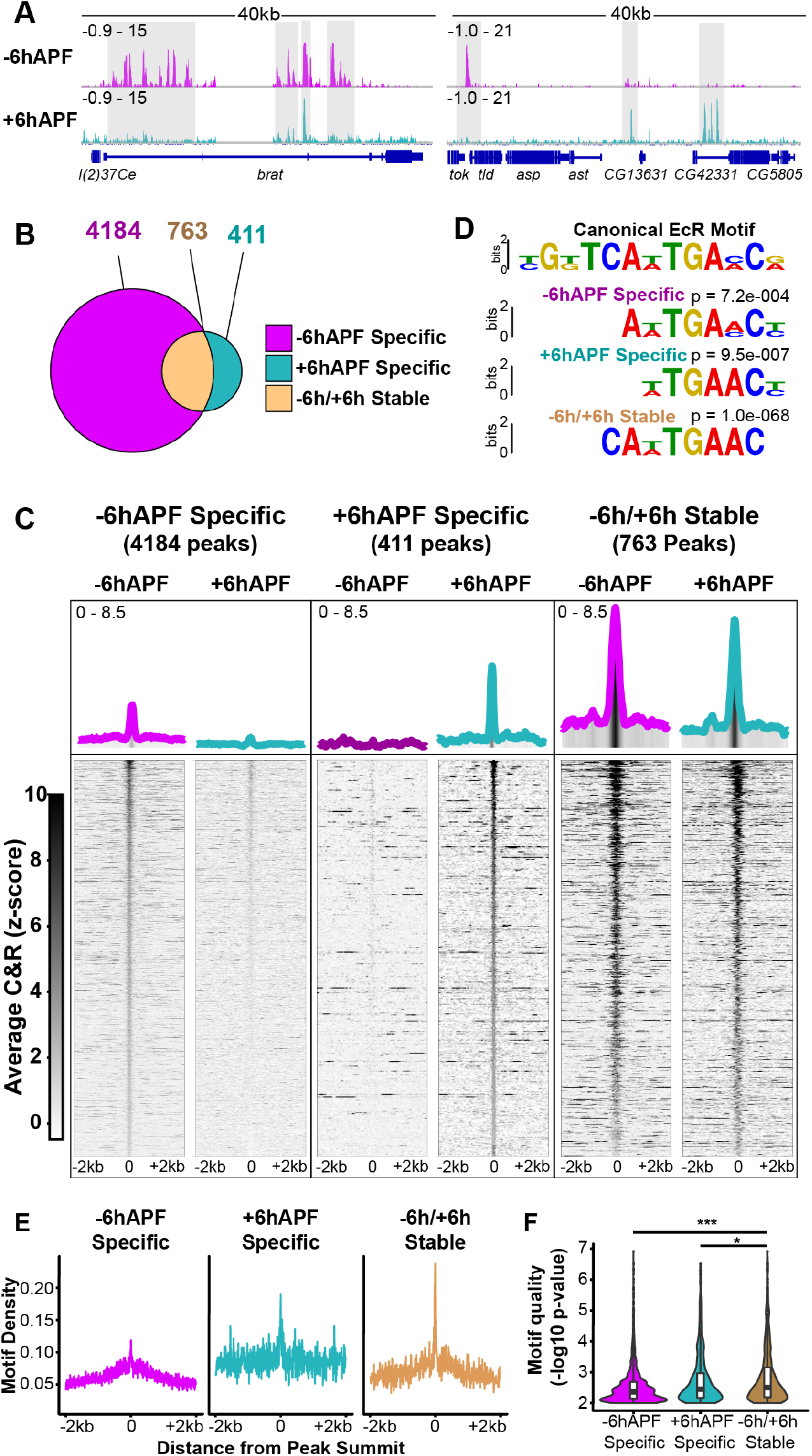
EcR binding is temporally dynamic. (A) Browser shots of EcR CUT&RUN data from −6hAPF and +6hAPF wings, with examples of −6hAPF unique, +6hAPF unique and −6h/+6h stable peaks highlighted by grey boxes. (B) Venn diagrams showing the number of peaks in each category. (D) Sequence logos comparing the canonical EcR/USP binding motif (top) to motifs identified through *de novo* motif analysis. (C) Heatmaps and average signal plots of EcR C&R signal (z-score). (E) Motif density plots of the number of EcR motifs around the peak summit. For –6h/+6h stable peaks, the motif summit for +6h was used. (F) Violin plots showing the average motif strength (–log_10_ p-value) of motifs within EcR peaks (* p-value < 0.05; *** p-value < 0.001, students t-test).

To investigate the potential biological significance of temporally-dynamic EcR binding, we separated EcR binding sites into three categories: –6hAPF-specific, +6hAPF-specific, and – 6h/+6h stable. Gene annotation enrichment analysis identified genes involved in imaginal disc-derived wing morphogenesis as the top term for each binding site category (**Table S3**), indicating that EcR may directly regulate genes involved in wing development at each of these developmental stages. Interestingly, we found that the amplitude of EcR CUT&RUN signal is greater at –6h/+6h stable binding sites relative to temporal-specific binding sites (**Fig 3C**). To investigate the potential basis for the difference in binding intensity, we examined the DNA sequence within each class of EcR binding site. Together with its DNA binding partner, Usp, EcR recognizes a canonical, 13bp palindromic motif, called an Ecdysone Response Element (EcRE), with EcR and Usp each binding to half of the motif (38, 39). *De novo* motif discovery analysis revealed the presence of this motif in each of our three peak categories (**Fig 3D**). To determine if differences in signal amplitude between –6h/+6h stable EcR binding sites could be caused by differences in motif content, we examined EcR motif density around the CUT&RUN peak summits for each of the three binding site categories. On average, we observed a positive correlation between motif density and signal amplitude, with –6hAPF temporal-specific binding sites having both the lowest motif density and the lowest signal amplitude, and –6h/+6h stable binding sites having both the highest motif density and the highest signal amplitude (**Fig 3E**). Furthermore, the average motif strength (ie. the extent to which the motif matches consensus) in –6h/+6h stable binding sites was also significantly higher (**Fig 3F**). These data are consistent with a model in which differences in motif content and strength within −6h/+6h stable peaks make EcR binding to these sites less reliant on other factors, such as other transcription factors. Conversely, the lower motif content and strength within temporal-specific peaks suggests EcR may depend more on cooperative interactions with other transcription factors to assist binding at these sites.

### EcR binding is tissue-specific

The results described above indicate that EcR binds extensively across the genome, including to many genes with wing-specific function, thus raising the question as to whether EcR binding is tissue-specific. To address this question, we first examined loci that had been previously determined to contain functional EcR binding sites by *in vitro* DNA binding and *in vivo* reporter assays (22, 38, 40). Many of these sites, including the glue genes *Sgs3, Sgs7,* and *Sgs8,* the fat body protein *Fbp1,* and the oxidative response gene *Eip71CD,* show no evidence of EcR binding in wings (**Fig S5**), supporting the finding that EcR binds target sites in a tissue-specific manner. To examine this question more globally, we compared our wing CUT&RUN data to EcR ChIP-seq data from *Drosophila* S2 cells (**Fig 4A**) (25). Overall, a small fraction of wing EcR binding sites overlap an EcR binding site in S2 cells (**Fig 4B, C**). However, among the sites that are shared between wings and S2 cells, there is marked enrichment of overlap with −6h/+6h stable wing binding sites. Whereas only 0.1% of −6hAPF-specific binding sites (41 peaks) and 2% of +6hAPF-specific binding sites (9 peaks) overlap an S2 cell EcR binding site, 16% of −6h/+6h stable binding sites (122 peaks) overlap an S2 cell EcR binding site. Thus, binding sites to which EcR is stably bound over time in developing wings are more likely to be shared with EcR binding sites in other cell types, relative to temporal-specific EcR binding sites in the wing.

**Figure 4:**
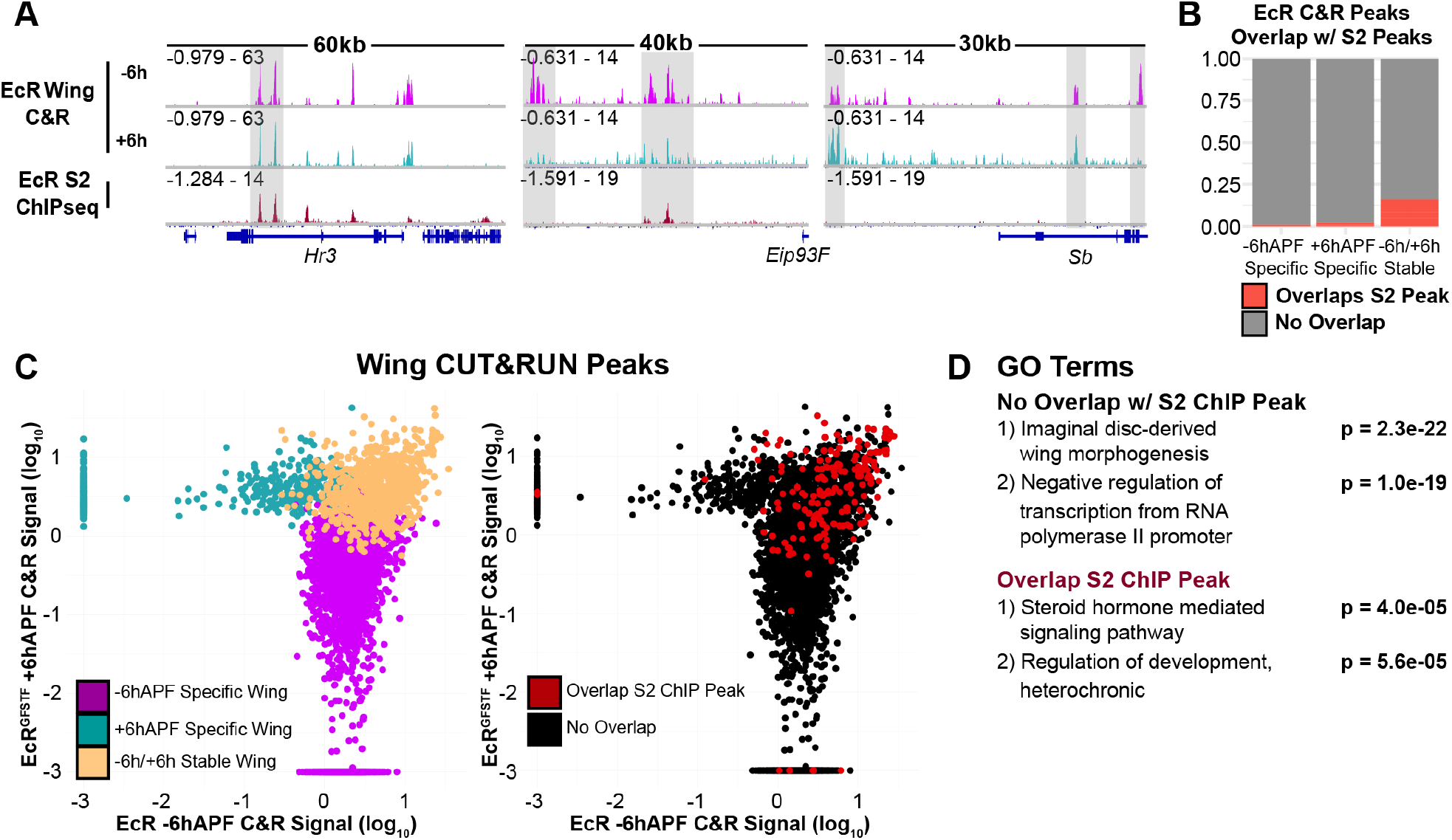
EcR binding is tissue-specific. (A) Browser shots comparing EcR CUT&RUN to EcR ChIPseq in S2 cells (25). Grey boxes highlight examples of shared (left), S2-specific (middle) and wing-specific peaks. (B) Bar plots showing the proportion of EcR C&R peaks that overlap an S2 ChIP peak in each category. (C) A comparison of the average signal within EcR C&R peaks colored by how they behave temporally (left) and whether they overlap an S2 ChIP peak (right). (D) GO terms of the closest gene to a wing EcR peak stratified by whether they overlap an S2 ChIP peak.

To investigate potential differences in target gene function between wing-specific binding sites and those shared with S2 cells, we performed gene annotation enrichment analysis on genes near EcR binding sites. This analysis revealed steroid hormone-mediated signaling pathway as the most significant term for genes overlapping an EcR peak in both wings and S2 cells (**Fig 4D**). Genes annotated with this term include canonical ecdysone-responsive genes, such as *Eip78C, Hr39* and *EcR* itself. By contrast, imaginal disc-derived wing morphogenesis was identified as the top term for genes near wing-specific EcR binding sites, similar to our findings from above. These data indicate that EcR binding sites that are shared by wings and S2 cells tend to occur at canonical ecdysone target genes, whereas wing-specific EcR binding sites tend to occur at genes with wing-specific functions. Together, these data suggest EcR plays a direct role in mediating the distinct gene expression responses to ecdysone exhibited by different cell types (41).

### EcR regulates the temporal activity of an enhancer for *broad,* a canonical ecdysone target gene

The results described above indicate that EcR binds to both canonical and non-canonical ecdysone target genes in the wing, and that EcR is required for temporal progression of wing transcriptional programs. We next sought to examine the relationship between EcR binding in the genome and regulation of gene expression. Because EcR both activates and represses target gene expression, we grouped all differentially expressed genes together and counted the proportion of genes that overlap an EcR binding cluster (**Fig S6A-C**). We observed an enrichment of EcR binding sites near genes that are differentially expressed in EcR-RNAi wing at both –6hAPF and +6hAPF and a depletion of EcR binding sites near genes that are either temporally static or not expressed (**Fig S6A-C**). These correlations support a direct role for EcR in regulating temporal changes in gene expression during the larval-to-prepupal transition.

To obtain a more direct readout of EcR’s role in target gene regulation, we investigated whether EcR binding contributes to control of enhancer activity. We first examined the potential regulation of a canonical ecdysone target gene. The *broad complex* (*br*) encodes a family of transcription factors that are required for the larval-to-prepupal transition in wings and other tissues (**Fig 5A**) (42, 43). *Br* has been characterized as a canonical ecdysone target gene that is induced early in the transcriptional response upon release of hormone (42, 44, 45). In wing imaginal discs, Br protein levels are uniformly low in early 3^rd^ instar larvae, and by late 3^rd^ instar, Br levels have increased (**Fig S7A**). Ecdysone signaling has been proposed to contribute to this increase in Br expression in wings over time (30, 31).

**Figure 5:**
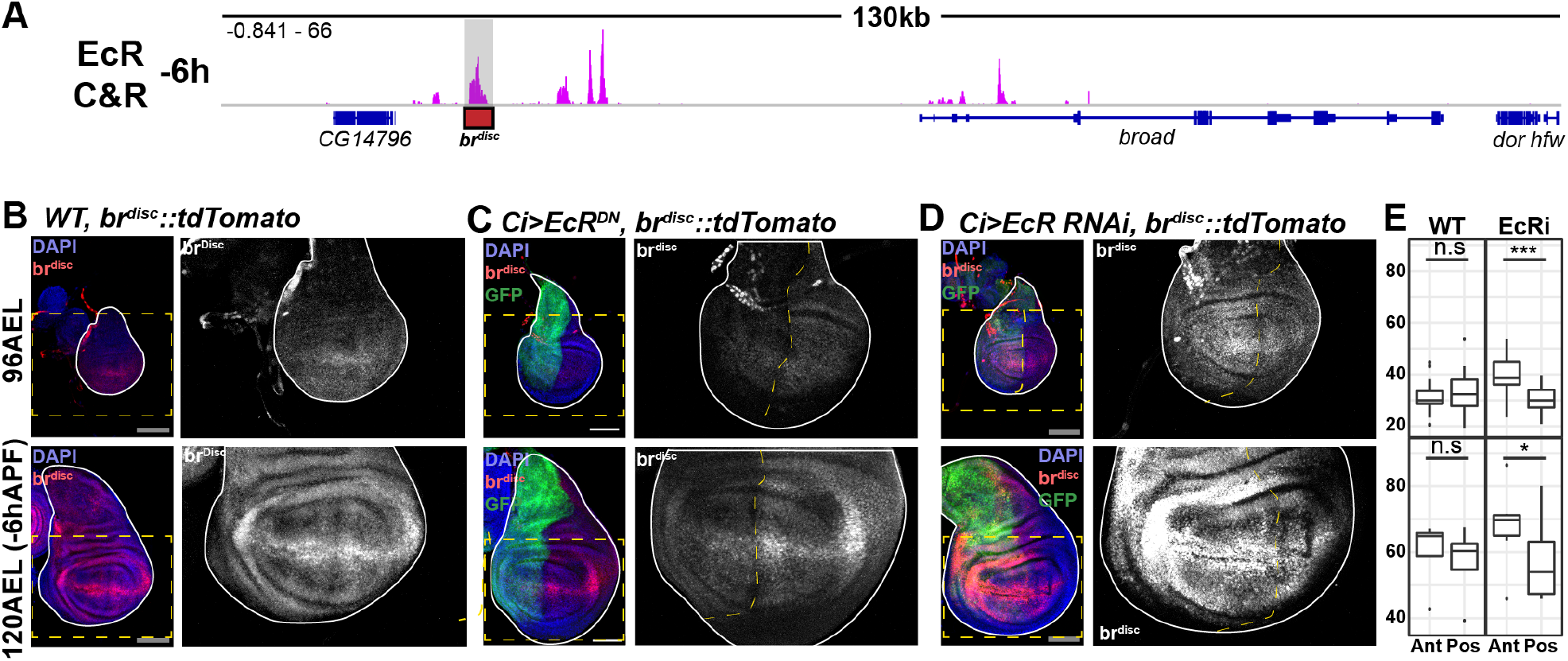
EcR regulates the temporal activity of an enhancer for the gene *broad*. (A) Browser shots of the *br* locus, with the location of the *br^disc^* highlighted by a shaded gray region. (B) *br^disc^* activity in WT wings (red) at 96hrs after egg laying (96AEL) and 120AEL (– 6hAPF). (C) The effect expressing EcR dominant negative (EcR^DN^) in the anterior compartment of the wing marked by GFP (green) on *br^disc^* activity. (D) Comparison of *br^disc^* activity between the anterior (Ant) and posterior (Pos) compartments of the wing in WT and EcR-RNAi wings (* p < 0.05; *** p < 0.005, paired student’s t-test). Dotted yellow boxes indicate the location of insets. Scale bars are 100uM.

Our CUT&RUN data identify multiple EcR binding sites across the *br* locus at both −6hAPF and +6hAPF (**Fig 5A**). One of these binding sites corresponds to an enhancer (*br^disc^*) we previously identified that recapitulates *br* activity in the wing epithelium at −6hAPF (26). Consistent with the observed increase in Br protein levels during 3^rd^ instar wing development, the activity of *br^disc^* increases with time (**Fig 5B**). To investigate the potential role of EcR in controlling the activity of *br^disc^*, we ectopically expressed a mutated isoform of EcR that functions as a constitutive repressor (EcR^DN^). EcR^DN^ expression in the anterior compartment of the wing results in decreased *br^disc^* activity in both early and late stage wing discs (**Fig 5C**), indicating that EcR^DN^ represses *br^disc^*. We further examined the role of EcR in regulating *br^disc^* by knocking down EcR via RNAi, which would eliminate both activating and repressing functions of EcR. EcR knockdown resulted in a modest increase in the activity of *br^disc^* in early wing discs compared to WT wings (**Fig 5D-E**), demonstrating that EcR is required to repress *br^disc^* at this stage. We also observed a slight increase in *br^disc^* activity in late wing discs (**Fig 5D-E**). Together, these findings indicate that EcR is required to keep *br^disc^* activity low in early 3^rd^ instar wing discs, but it is not required for *br^disc^* activation in late 3^rd^ instar wing discs. Additionally, the observation that *br^disc^* is active in the absence of EcR, and continues to increase in activity over time, suggests that *br* requires other unknown activators which themselves may be temporally dynamic. Because the levels of Br increase with time, we conclude that release of repression by EcR functions as a temporal switch to control Br expression during the larval-to-prepupal transition.

### EcR binds to enhancers with spatially-restricted activity patterns in the wing

EcR’s role in controlling the timing of *br* transcription through the *br^disc^* enhancer supports conventional models of ecdysone signaling in coordinating temporal gene expression. To determine whether EcR plays a similar role at non-canonical ecdysone target genes, we focused on the *Delta* (*Dl*) gene, which encodes the ligand for the Notch (N) receptor. Notch-Delta signaling is required for multiple cell fate decisions in the wing (46–48). In late third instar wing discs, *Dl* is expressed at high levels in cells adjacent to the dorsal-ventral boundary, along each of the four presumptive wing veins, and in proneural clusters throughout the wing (49). Remarkably, despite the requirement of Notch-Delta signaling in each of these areas, no enhancers active in wing discs have been described for the *Dl* gene. The *Dl* locus contains multiple sites of EcR binding (**Fig 6A**). Using open chromatin data from wing imaginal discs to identify potential *Dl* enhancers (50), we cloned two EcR-bound regions for use in transgenic reporter assays. The first of these enhancers exhibits a spatially-restricted activity pattern in late third instar wing discs that is highly reminiscent of sensory organ precursors (SOPs) (**Fig 6B**). Immunostaining for the proneural factor Achaete (Ac) revealed that cells in which this *Dl* enhancer is active co-localize with proneural clusters (**Fig 6B**). Immunostaining also confirmed these cells express Dl (**Fig 6C**). We therefore refer to this enhancer as *Dl^SOP^*. Notably, using *Dl^SOP^* to drive expression of a destabilized GFP reporter, its activity pattern refines from a cluster of cells to a single cell (**Fig 6C**), consistent with models of SOP specification in which feedback loops between *N* and *Dl* result in high levels of N signaling in the cells surrounding the SOP, and high levels of *Dl* expression in the SOP itself. By +6hAPF, the pattern of *Dl^SOP^* activity does not change, and it remains spatially restricted to cells along the D/V boundary and proneural clusters in the notum. The second *Dl* enhancer bound by EcR is also active in late 3^rd^ instar wing discs (**Fig 6A**). This enhancer is most strongly active in Dl-expressing cells of the tegula, lateral notum, and hinge (**Fig 6D-E**) (51). In the pouch, it is active in cells that comprise the L3 and L4 proveins, which require *Dl* for proper development (47) although overlap with Dl in each of these regions is less precise (**Fig 6D-E**). We refer to this enhancer as *Dl^teg^*. Collectively, these data demonstrate that, in contrast to the widespread activity of *br^disc^*, the EcR-bound enhancers in the *Dl* locus exhibit spatially-restricted activity, raising the possibility that EcR binding may serve a different function at these binding sites.

**Figure 6:**
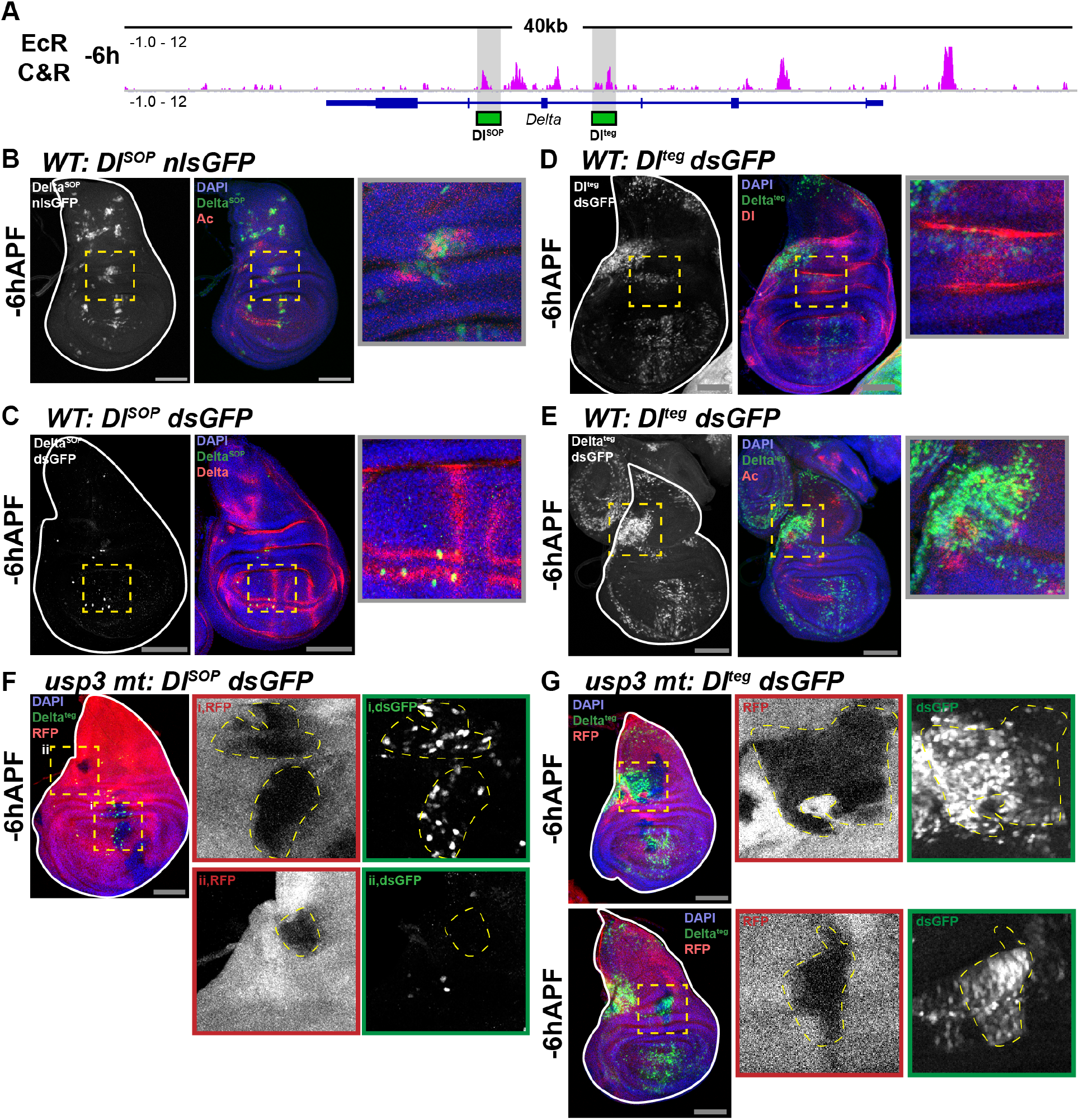
EcR regulates the spatial activity of enhancers for the gene *Dl*. (A) Browser shots of the *Dl* locus, with the location of the *Deltd^SOP^* and *Delt^F^* highlighted by a gray box. (B-C) Enhancer activity of *Dl^SOP^* (green) showing overlap with Ac and Dl. (D-E) Enhancer activity of *Dl^teg^* showing overlap with Dl and Ac. (F-G) Enhancer activity of *Dl^SOP^* and *Dl^teg^* in *usp^3^* mitotic clones which are marked by the absence of RFP. Dotted yellow boxes indicate the location of insets. Scale bars are 100μm.

### *Ultraspiracle* clones display changes in the spatial pattern of enhancer activity

We next sought to determine if EcR regulates the activity of these enhancers. Since the *Dl* enhancers drive GAL4 expression, we could not use the EcR^DN^ and EcR-RNAi lines employed above. Therefore, we generated loss of function clones of Usp, the DNA binding partner of EcR. Clones of *usp* were induced at 48-60 hours and enhancer activity was assayed at −6hAPF. Surprisingly, *usp* loss of function results in an increased number of cells in which *Dl^SOP^* is active in the pouch of wing discs (**Fig 6F, inset i**), suggesting that EcR/Usp are required to repress *Dl^SOP^* activation. Notably, clones of *usp* in other regions of the wing (**Fig 6F, inset ii**) do not activate *Dl^SOP^,* indicating that EcR/Usp are not necessary for repression of *Dl^SOP^* in all cells of the wing. We also note that regions exhibiting ectopic *Dl^SOP^* activity in *usp* clones tend to be near regions of existing *Dl^SOP^* activity, suggesting that localized activating inputs are required to switch the *Dl^SOP^* enhancer on, and that EcR/Usp binding to *Dl^SOP^* acts as a countervailing force to restrict its activation to certain cells within these regions. Because the pattern of *Dl^SOP^* activity does not expand between –6hAPF and +6hAPF in WT wings, the ectopic activation of this enhancer in *usp* clones supports the conclusion that EcR/Usp regulate the spatial pattern of *Dl^SOP^* activation rather than its temporal activity pattern, as in the case of the *br^disc^* enhancer.

We observed a similar effect of *usp* loss of function on activity of the *Dl^teg^* enhancer. *Dl^teg^* activity expands in *usp* clones adjacent to regions in which *Dl^teg^* is active in WT cells (**Fig 6F**). As with *Dl^SOP^,* however, loss of *usp* function does not appear to be sufficient to cause ectopic *Dl^teg^* activity, as clones that are not adjacent to existing *Dl^teg^* activity do not ectopically activate the enhancer. Notably, we did not observe expanded expression of Ac within *usp* clones, suggesting that the expanded activity pattern of the clones is not due to an expanded proneural domain (**Fig S8**). These results suggest that EcR primarily functions to repress these enhancers at –6hAPF in order to spatially restrict their activity. The observation that *usp* loss of function is not sufficient to cause ectopic enhancer activity may be because the activation of these enhancers requires other inputs.

## DISCUSSION

Decades of work have established the central role that ecdysone signaling, acting through its nuclear receptor, EcR, plays in promoting developmental transitions in insects. In this study, we investigate the genome-wide role of EcR during the larval-to-prepupal transition in *Drosophila* wings. Our findings validate existing models of ecdysone pathway function, and they extend understanding of the direct role played by EcR in coordinating dynamic gene expression programs.

### The role of EcR in promoting gene expression changes during developmental transitions

Our RNA-seq data reveal that EcR controls the larval-to-prepual transition by activating and repressing distinct sets of target genes. In larval wing imaginal discs, we find that EcR is primarily required to prevent precocious activation of the prepupal gene expression program. This finding is consistent with previous work which demonstrated precocious differentiation of sensory neurons in the absence of ecdysone receptor function (33). Since ecdysone titers remain low during most of the 3^rd^ larval instar, these data are also consistent with prior work which demonstrated that EcR functions as a transcriptional repressor in the absence of hormone (6, 38). Later in prepupal wings, we find that EcR loss of function results in failure to activate the prepupal gene expression program. Indeed, many of the genes that become precociously activated in wing discs fail to reach their maximum level in prepupae. Since rising ecdysone titers at the end of 3^rd^ larval instar trigger the transition to the prepupal stage, this finding is consistent with a hormone-induced switch in EcR from a repressor to an activator (6, 38). We also find that EcR loss of function results in persistent activation of the larval gene expression program in prepupal wings. This finding is not clearly explained by a hormone-induced switch in EcR’s regulatory activity. However, it is possible that EcR activates a downstream transcription factor, which then represses genes involved in larval wing development. Overall, these findings indicate that EcR functions both as a temporal gate to ensure accurate timing of the larval-to-prepupal transition and as a temporal switch to simultaneously shut down the preceding developmental program and initiate the subsequent program. Finally, it is of particular note that these genome-wide results fit remarkably well with the model of ecdysone pathway function predicted by Ashburner forty-five years ago (52).

### Widespread binding of EcR across the genome

Existing models describe EcR as functioning at the top of a transcriptional cascade, in which it binds to a relatively small number of canonical ecdysone-response genes. These factors then activate the many downstream effectors that mediate the physiological response to ecdysone. Consistent with this model, attempts to assay EcR binding genome-wide in S2 cells and Kc167 cells have identified relatively few EcR binding sites (24, 25). However, this model does not adequately explain how ecdysone elicits distinct transcriptional responses from different target tissues. Our data reveal that EcR binds to thousands of sites genome-wide. While many of the genes that exhibit EcR binding sites have previously been identified as direct targets of EcR in salivary glands and others tissues, the majority of EcR binding events we observe occur near genes with essential roles in wing development. These data support a model in which EcR directly mediates the response to ecdysone both at the top of the hierarchy and at many of the downstream effectors. Interestingly, comparison of our wing DNA-binding profiles with ChIP-seq data from S2 cells revealed that shared EcR binding sites are enriched in canonical ecdysone-response genes, suggesting that the top tier of genes in the ecdysone hierarchy are direct targets of EcR across multiple tissues, while the downstream effectors are direct EcR targets only in specific tissues. These data neatly account for the observation that parts of the canonical ecdysone transcriptional response are shared between tissues, even as many other responses are tissue-specific. It will be important to identify the factors that contribute to EcR’s tissue-specific DNA targeting in future work. It is possible that tissue-specific transcription factors facilitate EcR binding across the genome, as suggested by recent DNA-binding motif analysis of ecdysone-responsive enhancers in S2 and OSC cell lines (25). Alternatively, tissue-specific epigenetic marks such as histone modifications may influence EcR binding to DNA.

### Temporally-dynamic binding of EcR

Pulses of ecdysone mediate different transcriptional responses at different times in development. Some of this temporal-specificity has been shown to be mediated by the sequential activation of transcription factors that form the core of the ecdysone cascade (53–55). Our data suggest that changes in EcR binding may also be involved. We find that EcR binding is highly dynamic over time; a subset of its binding sites is unique to each time point. The mechanisms responsible for changes in EcR binding over time remain unclear. One possibility is that ecdysone titers impact EcR DNA binding. This could occur through ligand-dependent changes in EcR structure or through ligand-dependent interactions with co-activator and co-repressor proteins that influence EcR’s DNA binding properties. An alternative possibility is that the nuclear-to-cytoplasmic ratio of EcR changes with time, as has been previously proposed (56, 57). While nuclear export of EcR could explain the global reduction in the number of EcR binding sites between –6hAPF and +6hAPF, it cannot explain the appearance of new EcR binding sites at +6hAPF. For this reason, it is notable that temporal-specific binding sites contain lower motif content on average relative to EcR binding sites that are stable between –6hAPF and +6hAPF. This suggests that temporal-specific binding may be more dependent on external factors. An intriguing possibility is that stage-specific transcription factors activated as part of the canonical ecdysone cascade may contribute to recruitment or inhibition of EcR binding at temporal-specific sites.

### EcR controls both temporal and spatial patterns of gene expression

EcR has been shown to act as both a transcriptional activator and a repressor. This dual functionality confounded our attempts to draw genome-wide correlations between EcR binding and changes in gene expression. Therefore, we sought to determine the effect of EcR binding by examining individual target enhancers. We find that EcR regulates the temporal activity of an enhancer for the canonical early-response target gene, *br.* In wild type wings, the activity of this enhancer increases between early and late third instar stages, as do Br protein levels. Ectopic expression of a dominant-repressor isoform of EcR decreased activity of *br^disc^*. Surprisingly, RNAi-mediated knockdown of EcR increased *br^disc^* activity, indicating that EcR is not required for *br^disc^* activation. Instead, these findings indicate that EcR represses *br^disc^* in early third instar wings, consistent with our RNA-seq data which demonstrated that EcR prevents precocious activation of the prepupal gene expression program prior to the developmental transition. It is not known what factors activate *br* or the other prepupal genes.

Temporal control of gene expression by EcR is expected given its role in governing developmental transitions. However, our examination of EcR-bound enhancers from the *Dl* locus demonstrates that it also directly controls spatial patterns of gene expression. Loss-of-function clones for EcR’s DNA binding partner Usp exhibited ectopic activation of two *Dl* enhancers. However, we did not detect ectopic enhancer activity in all *usp* mutant clones, indicating that EcR is required to restrict activity of target enhancers only at certain locations within the wing. Examination of +6hAPF wings revealed no changes in the spatial pattern of *Dl* enhancer activity relative to −6hAPF, indicating that ectopic enhancer activation in *usp* clones does not reflect incipient changes in enhancer activity. Recently, EcR binding sites were shown to overlap with those for the *Notch* regulator, Hairless, supporting a potential role of EcR in regulating spatial patterns of gene expression. (59). We conclude that EcR regulates both temporal and spatial patterns of gene expression. Given the widespread binding of EcR across the genome, our findings suggest that EcR plays a direct role in temporal and spatial patterning of many genes in development.

Hormones and other small molecules act through nuclear receptors to initiate transcriptional cascades that often continue for extended periods of time. For example, thyroid hormone triggers metamorphosis in frogs and other chordates, a process that can take weeks for completion (60). Our work raises the possibility that nuclear receptors play an extensive and direct role in regulating activity of downstream response genes. In particular, the widespread and temporally-dynamic binding of EcR that we observed over a short interval of wing development suggests that the complete repertoire of EcR targets is vastly larger than previously appreciated.

## METHODS

### Western Blots

For each sample, 40 wings were lysed directly in Laemmlli sample buffer preheated to 95C. The following antibody concentrations were used to probe blots: 1:1000 mouse anti-EcR (DSHB DDA2.7, concentrate); 1:5000 rabbit anti-GFP (Abcam ab290); 1:30000 mouse anti-alpha Tubulin (Sigma T6074); 1:5000 goat anti-mouse IgG, HRP-conjugated (Fisher 31430); 1:5000 donkey anti-rabbit, HRP-conjugated (GE Healthcare NA934).

### Transgenic Reporter Construction

Candidate enhancers were cloned into the pΦUGG destination vector (61) <u>and</u> integrated into the attP2 site. Primer sequences are available upon request.

### Immunofluorescence

Immunostaining was performed as described previously (50). For mitotic clones, *usp3 FRT19A / Ubi-RFP, hs-FLP, FRT19A; Enhancer-GAL4 / UAS-dsGFP* animals were heat-shocked at 24-48hrs AEL. The following antibody concentrations were used: 1:750 mouse anti-EcR, 1:4000 rabbit anti-GFP, 1:3500 mouse anti-Dl (DSHB C594.9b, concentrate), 1:200 mouse anti-FLAG M2 (Sigma F1804), 1:10 mouse anti-Achaete (DSHB anti-achaete, supernatant).

### Sample preparation for RNAseq

A minimum of 60 wings were prepared as previously described (McKay and Lieb, 2013) from either *Oregon R* (WT) or *yw; vg-GAL4, tub>CD2>GAL4, UAS-GFP, UAS-FLP / UAS-EcR-RNAi^104^* (EcR-RNAi). For library construction, 50-100ng RNA was used as input to the Ovation *Drosophila* RNA-Seq System. Single-end, 1×50 sequencing was performed on an Illumina HiSeq 2500 at the UNC High Throughput Sequencing Facility.

### Sample preparation for CUT&RUN

A minimum of 100 wings from *w; EcR^GFSTF^/Df(2R)BSC313* were dissected in 1XPBS. Samples were centrifuged at 800rcf for 5minutes at 4C and washed twice with dig-wash buffer (20mM HEPES-NaOH, 150mM NaCl, 2mM EDTA, 0.5mM Spermidine, 10mM PMSF, 0.05% digitonin) and incubated in primary antibody for 2hrs at 4C. Samples were washed as before and incubated in secondary antibody for 2hrs. Samples were washed and incubated for 1hr with proteinA MNase. Samples were washed twice in dig-wash buffer without EDTA and then resuspended in 150uL dig-wash buffer without EDTA. Following this, samples were equilibrated to 0C in an ice bath. 2uL CaCl_2_ (100mM) was added to activate MNase and digestion allowed to proceed for 45s before treating with 150uL 2XRSTOP+ buffer (200mM NaCl, 20mM EDTA, 4mM EGTA, 50ug/ml RNase, 40ug/ml glycogen, 2pg/ml yeast spike-in DNA). Soluble fragments were released by incubating at 37C for 10m. Samples were spun twice at 800g, 5m at 4C and the aqueous phase removed. The rest of the protocol was performed as described in Skene et al., 2018. For library preparation, the Rubicon Thruplex 12s DNA-seq kit was used following the manufacturer’s protocol until the amplification step. For amplification, after the addition of indexes, 16-21 cycles of 98C, 20s; 67C, 10s were run. A 1.2x SPRI bead cleanup was performed (Agencourt Ampure XP). Libraries were sequenced on an Illumina MiSeq. The following antibody concentrations were used: 1:300 mouse anti-FLAG M2; 1:200 rabbit anti-Mouse (Abcam ab46450); 1:400 Batch#6 proteinA-MNase (from Steven Henikoff).

### RNA Sequencing Analysis

Reads were aligned with STAR (2.5.1b) (62). Indexes for STAR were generated with parameter –sjdbOverhang 49 using genome files for the dm3 reference genome. The STAR aligner was run with parameters –alignIntronMax 50000 –alignMatesGapMax 50000. Subread (v1.24.2) was used to count reads mapping to features (63). DESeq2 (v1.14.1) was used to identify differentially expressed genes using the lfcShrink function to shrink log-fold changes (64). Differentially expressed genes were defined as genes with an adjusted p-value less than 0.05 and a log2 fold change greater than 2. Normalized counts were generated using the counts function in DESeq2. For k-medoids clustering, normalized counts were first converted into the fraction of maximum WT counts and clustering was performed using the cluster package in R. Optimal cluster number was determined by minimizing the cluster silhouette. Heatmaps were generated using pheatmap (v1.0.10) in R. Gene Ontology analysis was performed using Bioconductor packages TopGO (v2.26.0) and GenomicFeatures (v1.26.4) (65, 66).

### CUT&RUN Sequencing Analysis

Technical replicates were merged by concatenating fastq files. Reads were trimmed using bbmap (v37.50) with parameters ktrim=4 ref=adapters rcomp=t tpe=t tbo=t hdist=1 mink=11. Trimmed reads were aligned to the dm3 reference genome using Bowtie2 (v2.2.8) with parameters –local –very-sensitive-local –no-unal –no-mixed –no-discordant –phred33 −I 10 −X 700 (67). Reads with a quality score less than 5 were removed with samtools (v1.3.1) (68). PCR duplicates were marked with Picard (v2.2.4) and then removed with samtools. Bam files were converted to bed files with bedtools (v2.25.0) with parameter -bedpe and split into different fragment size categories using awk (69). Bedgraphs were generated with bedtools and then converted into bigwigs with ucsctools (v320) (70). Data was z-normalized using a custom R script. MACS (v2016-02-15) was used to call peaks on individual replicates and merged files using a control genomic DNA file from sonicated genomic DNA using parameters −g 121400000 –nomodel –seed 123 (71). A final peak set was obtained by using peaks that were called in the merged file that overlapped with a peak called in at least one replicate. Heatmaps and average signal plots were generated from z-normalized data using the Bioconductor package Seqplots (v1.18.0). ChIPpeakAnno (v3.14.0) was used to calculate distance of peaks to their nearest gene (72, 73). Gene ontology analysis was performed as described above.

### Motif Analysis

*De novo* motif analysis was performed using DREME (v4.12.0) using parameters -maxk 13 −t 18000 −e 0.05 (74). As background sequences, FAIRE peaks from −6hAPF or +6hAPF were used. To identify occurrences of the EcR motif in the genome, a PWM for the canonical EcR motif was generated with the iupac2meme tool using the IUPAC motif “RGKTCAWTGAMCY” and then FIMO (v4.12.0) was run on the dm3 reference genome using parameters –max-stored-scores 10000000 –max-strand –no-qvalue –parse-genomic-coord –verbosity 4 –thresh 0.01 (75). Motif density plots were generated by counting the number of motifs from peak summits (10bp bins) and normalizing by the number of input peaks.

### Drosophila culture and genetics

Flies were grown at 25C under standard culture conditions. Late wandering larvae were used as the −6hAPF timepoint. White prepupae were used as the 0h time point for staging +6hAPF animals. For 96hAEL, apple juice plates with embyos were cleared of any larvae and then four hours later any animals that had hatched were transferred to vials. The following genotypes were used:

*yw; vg-GAL4, UAS-FLP, UAS-GFP, Tub>CD2>GAL4 / CyO (76)*
*w^1118^; P{UAS-EcR-RNAi}104* (BDSC#9327)
*yw; EcR^GFSTF^* (BDSC#59823)
*w1118; Df(2R)BSC313/CyO* (BDSC#24339)
*UAS-dsGFP* (gift of Brian McCabe)
*usp3, w*, P{neoFRT} 19A/FM7c* (BDSC#64295)
*P{Ubi-mRFP.nls}1, w*, P{hsFLP}12 P{neoFRT}19A* (BDSC#31418)

## ACKNOWLEDGEMENTS

We thank Peter J. Skene and Steven Henikoff for reagents and advice on the CUT&RUN protocol. Stocks obtained from the Bloomington Drosophila Stock Center (NIH P40OD018537) were used in this study. CMU was supported in part by NIH grant T32GM007092. This work was supported in part by Research Scholar Grant RSG-17-164-01-DDC to DJM from the American Cancer Society, and in part by grant R35GM128851 to DJM from the National Institute of General Medical Sciences of the NIH (https://www.nigms.nih.gov/).

## Supplemental Information

**Fig. S1.**
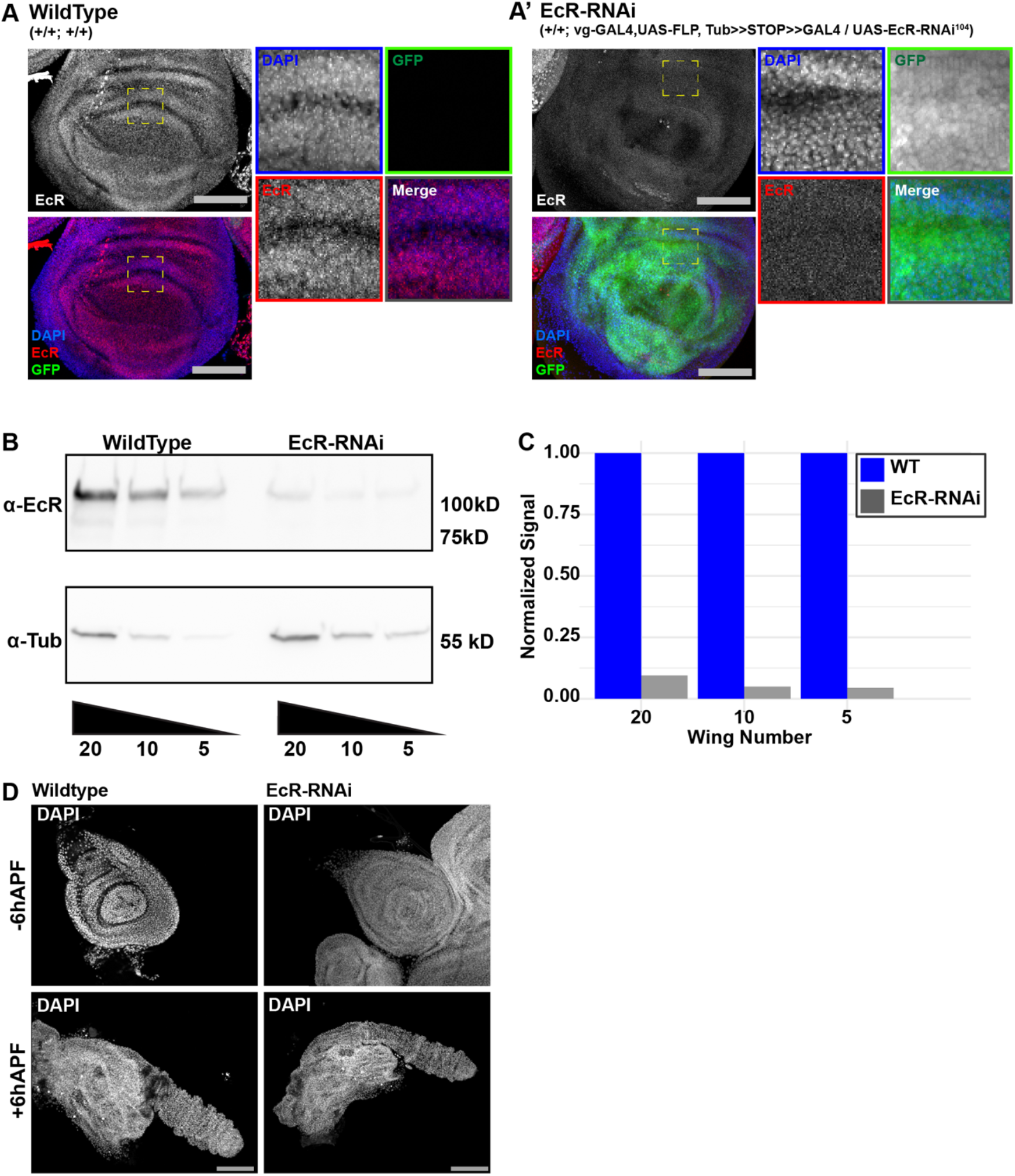
EcR-RNAi knock down is effective and does not result in systemic developmental arrest. (A) WT and *vg, tub-GAL4, UAS-EcR-RNAi* (hereafter EcR-RNAi) wings at –6hAPF. Location of insets is indicated by dashed boxes. (B) Western blots of EcR and alpha-tubulin levels in WT and EcR-RNAi wings from a serial dilution of wing tissue. (C) Quantification of western blots normalized to alpha-tubulin expressed as the fraction of WT signal. (D) Legs from WT and EcR-RNAi legs at −6hAPF and +6hAPF. Scale bars are 100μm.

**Fig. S2.**
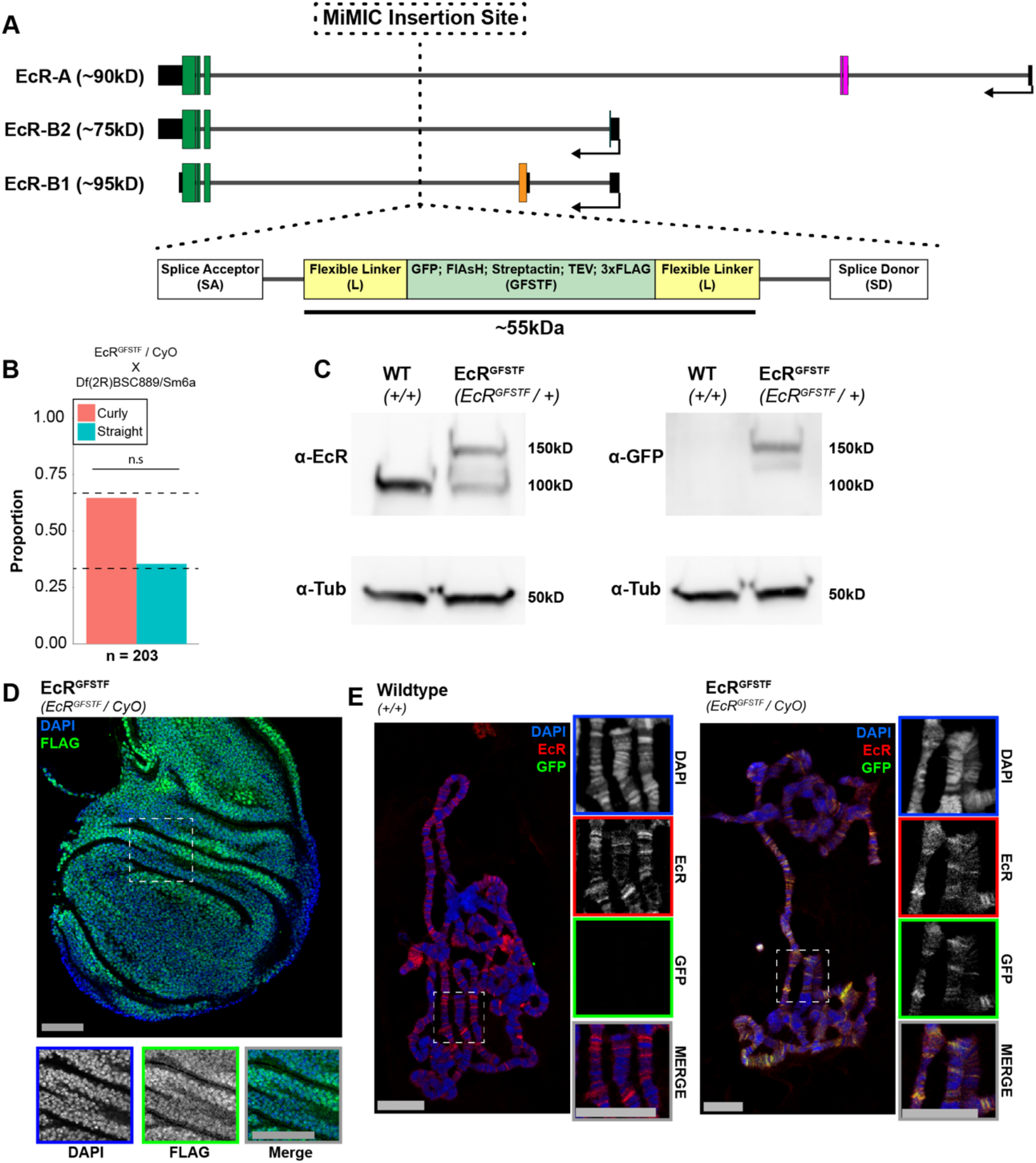
The EcR^GFSTF^ tag does not impair EcR function. (A) Location of the MiMIC insertion point in the EcR locus. The structure and size of the tag is indicated below. The insertion point is upstream of all exons shared between EcR isoforms and downstream of all isoform-specific exons. (B) Viability assay of *EcR^GFSTF^* animals crossed to a deficiency spanning the EcR locus. Statistical significance was determined using a chi-squared test with an expected ratio of 1:2 homozygous to heterozygous animals. (C) Western blots of wings from *EcR^GFSTF^* or WT animals stained for EcR or EcR^GFSTF^ (anti-GFP). (D) Immunostaining for *EcR^GFSTF^* (anti-FLAG) shows nuclear localization in wings. Scale bars are 50μm (E) Polytene squashes from WT or *EcR^GFSTF^*. Scale bars are 25μm. Dashed boxes indicate the location of insets.

**Fig. S3.**
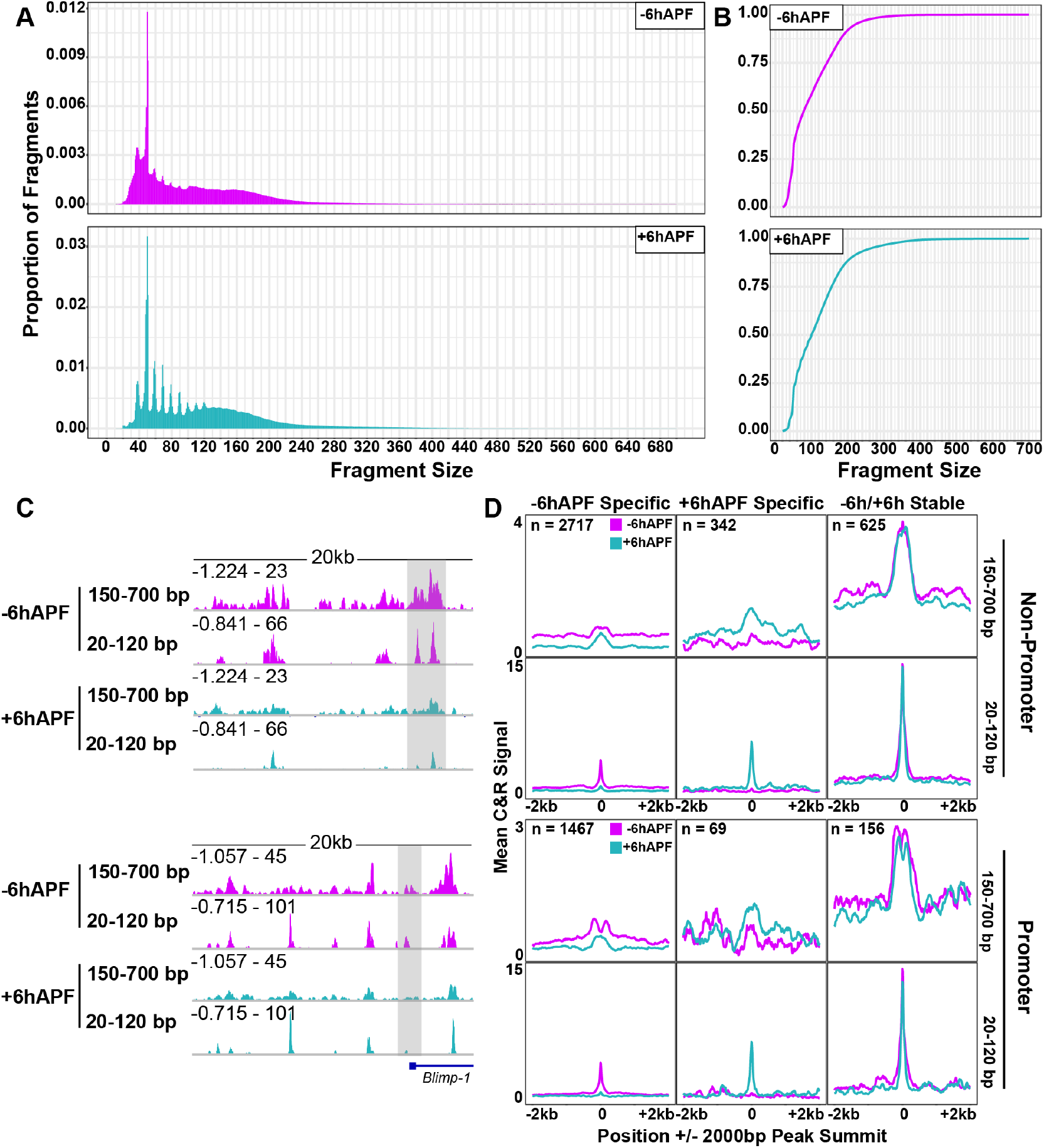
EcR CUT&RUN exhibits similar properties to those that have been previously reported. (A) Histograms of fragment sizes from EcR CUT&RUN. (B) Cumulative distribution plot of fragment sizes. (C) Representative browser shots comparing EcR C&R signal from 20-120 bp fragments and 150-700 bp. (D) Average signal plots of EcR C&R signal split by overlap with annotated promoters and fragment size.

**Fig. S4.**
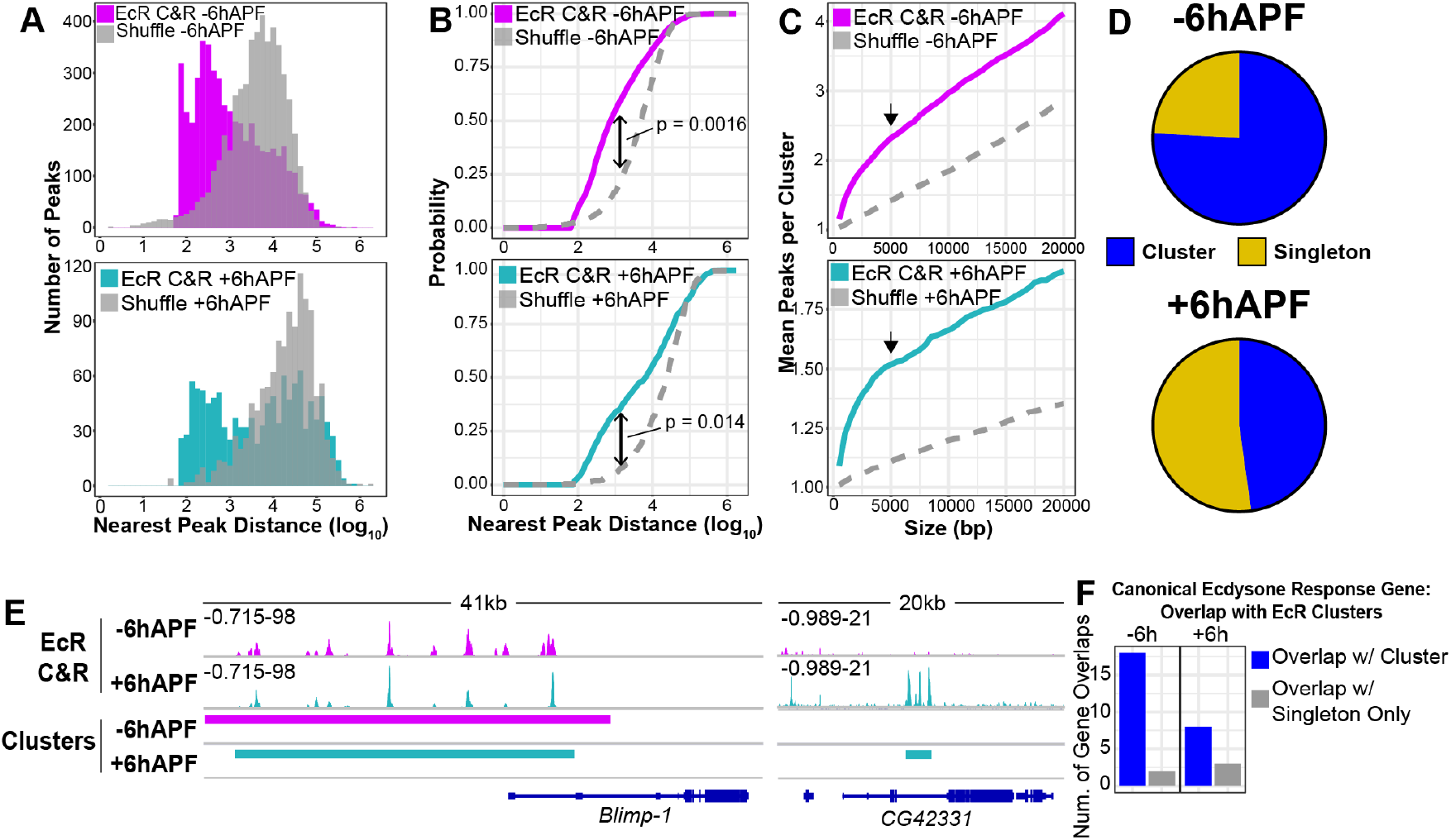
EcR peaks are clustered genome-wide. (A) Histograms of distance of each EcR peak to its nearest neighbor compared to a peak set shuffled over FAIRE peaks. (B) Cumulative distribution plots of the distance of each EcR peak to its nearest neighbor compared to shuffled peaks. Distributions were compared with a KS-test. (C) The mean number of peaks that overlap at least one other peak using different sizes of EcR peak. 5000bp (arrow) was used to define clusters in subsequent analyses. (D) Numbers of EcR peaks that fall into a cluster at −6hAPF and +6hAPF. (E) Examples of EcR clusters. (F) Numbers of canonical ecdysone-response genes that overlap an EcR cluster compared to those that only overlap an EcR singleton (ie. non-clustered peak). Canonical ecdysone response genes were defined as the union set of genes (42 total) in gene ontology terms: Cellular response to ecdysone (GO:0071390); Steroid hormone mediated signaling pathway (GO:0043401).

**Fig. S5.**
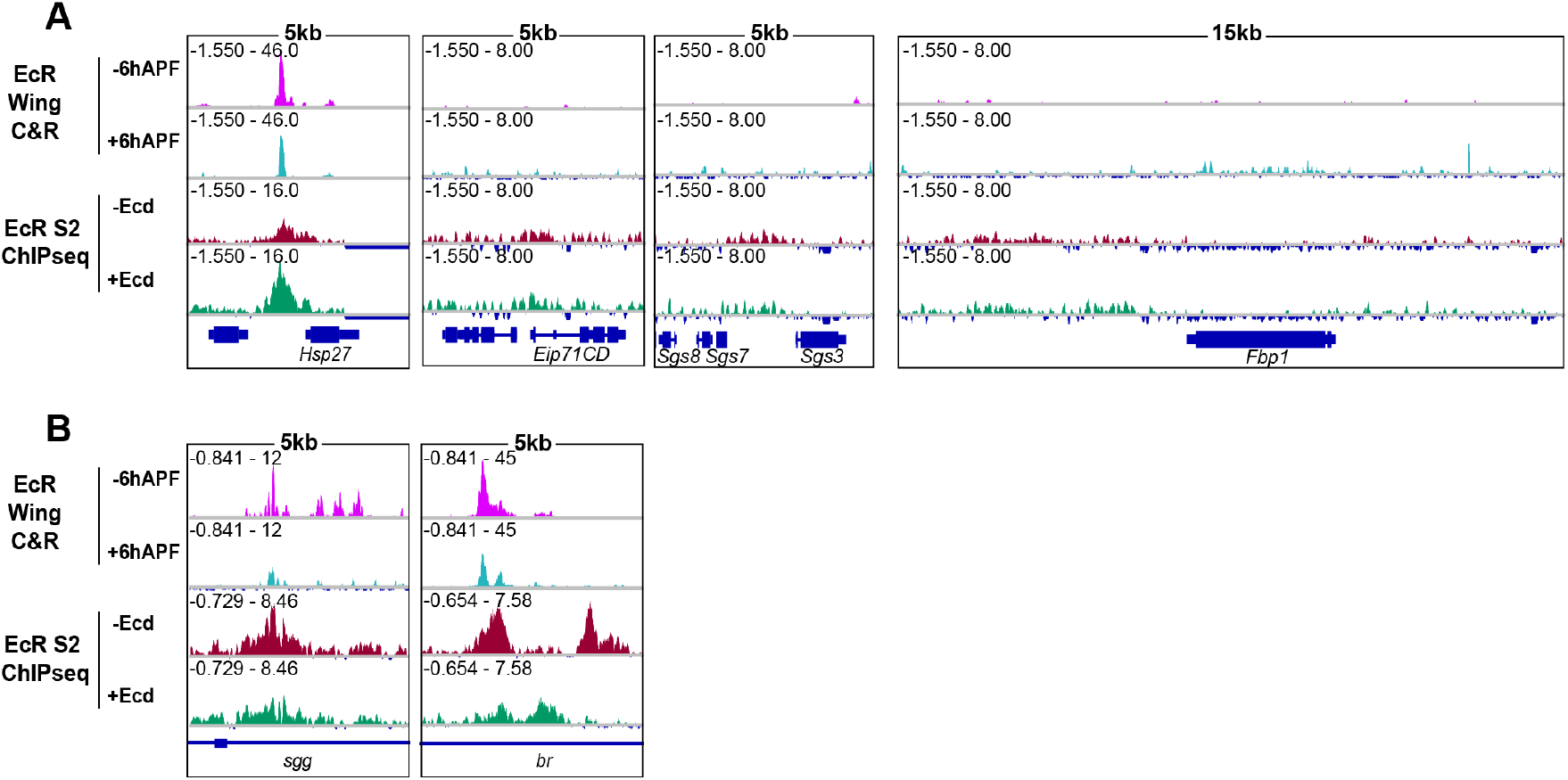
EcR binding is absent in wings and S2 cells from many sites previously identified as functional EcR binding sites in other tissues. (A) Browser shots showing EcR C&R signal and S2 ChIPseq (1) at previously identified EcR binding sites. (B) Browser shots comparing precision of EcR binding between EcR and S2 cells.

**Fig. S6.**
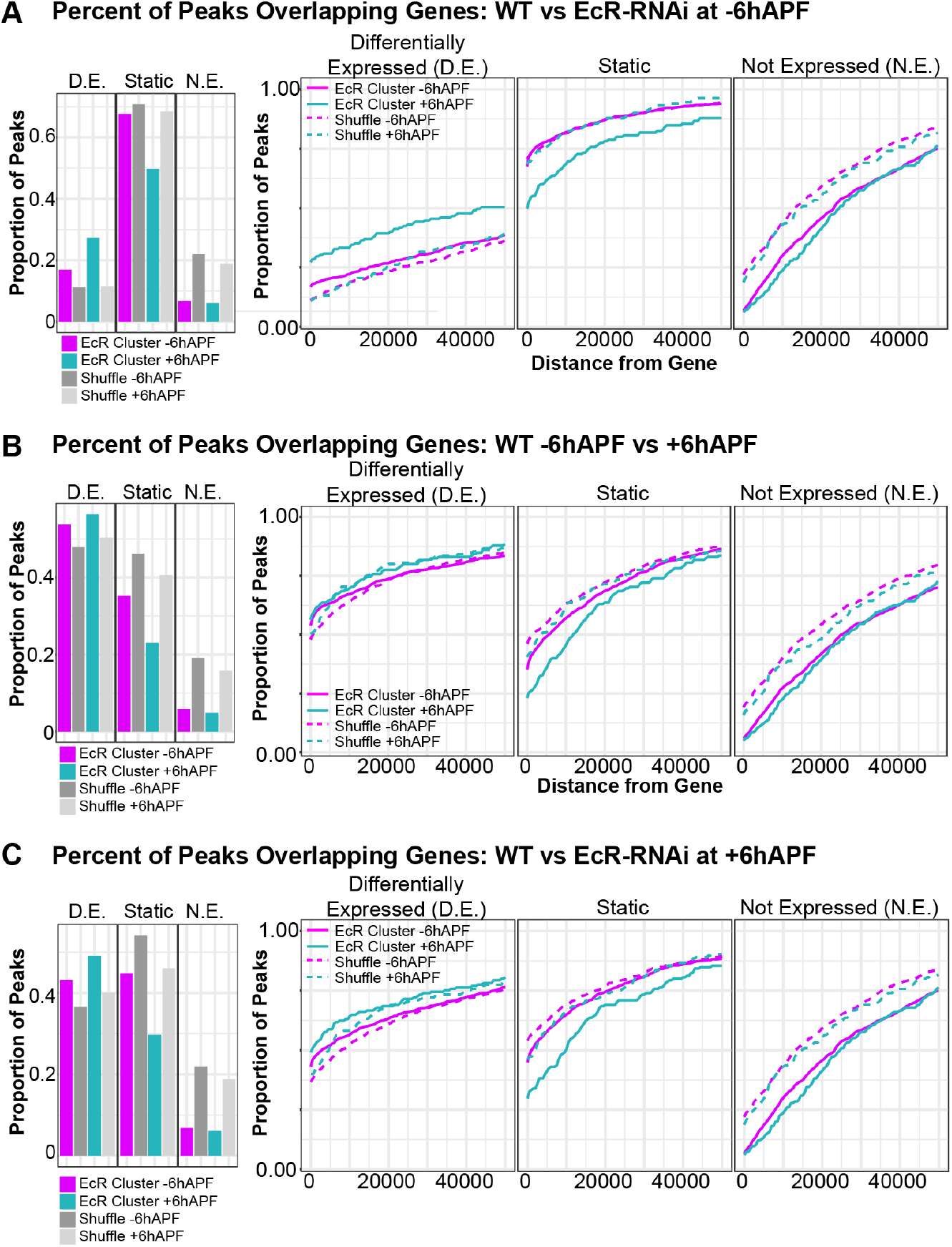
EcR binding is enriched at genes that are affected by EcR knockdown. Percentage of EcR clusters that overlap (left), or fall within some distance of (right), a differentially expressed (D.E.), static, or not-expressed (N.E.) gene in RNAseq comparing (A) WT to EcR-RNAi wings at −6hAPF, (B) WT −6hAPF to +6hAPF, (C) WT to EcR-RNAi at +6hAPF. EcR peaks were compared to peaks randomly shuffled over FAIRE peaks. Differentially expressed genes were defined as genes with an adjusted p-value < 0.05. Not expressed genes were defined as genes that were filtered out by DESeq2 (padj = NA).

**Fig. S7.**
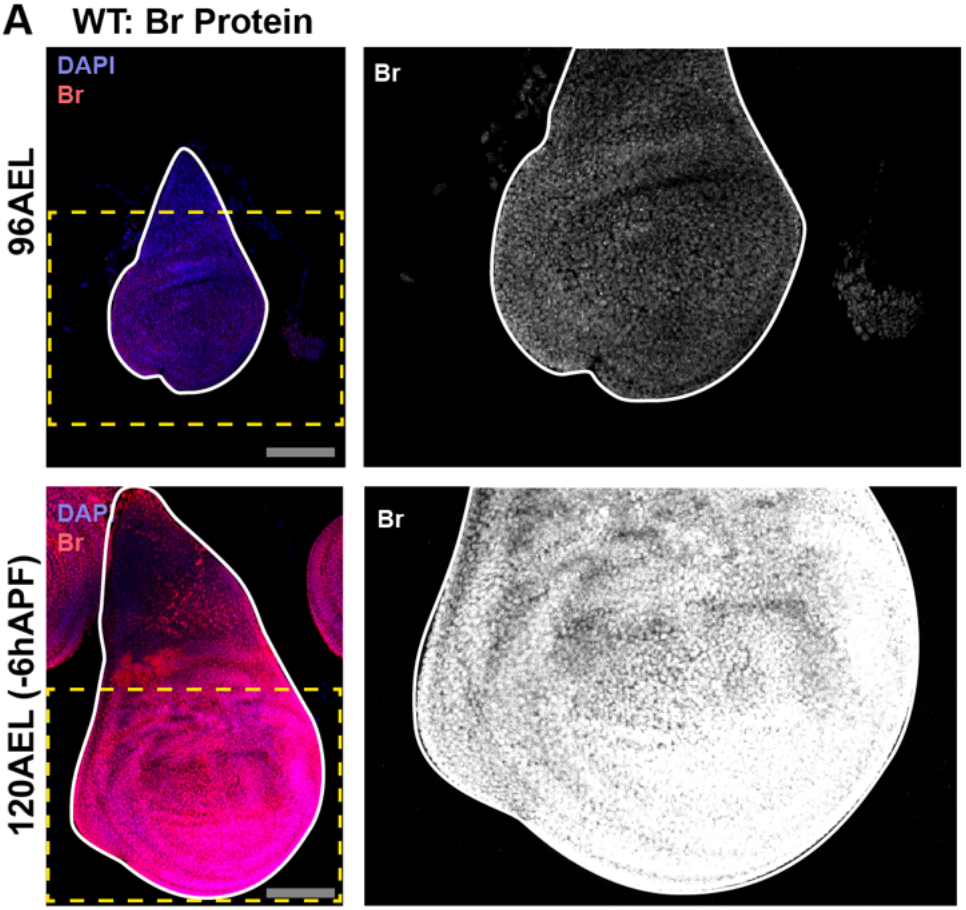
Broad protein levels increase with time. (A) Changes in Br protein (red) levels over time in WT wings between 96hrs after egg laying (96AEL) and 120AEL (-6hAPF). Scale bars are 100um. DAPI was used to stain nuclei.

**Fig. S8.**
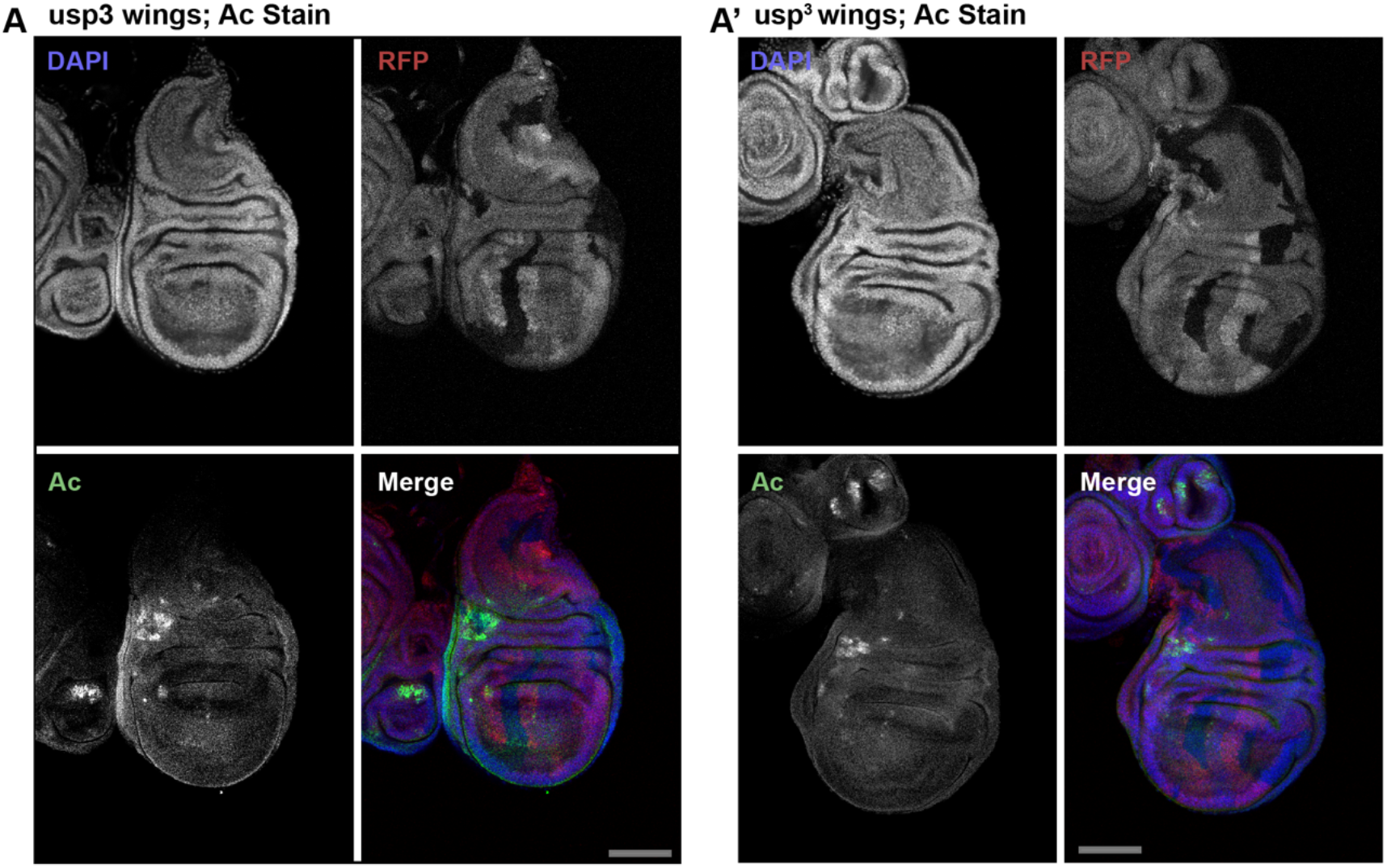
usp^3^ clones to not result in cell fate changes. (A) −6hAPF wings showing usp^3^ mitotic clones stained for Ac. Clones are marked by the absence of RFP. Scale bars are 100um. DAPI was used to stain nuclei.

**Table S1:**
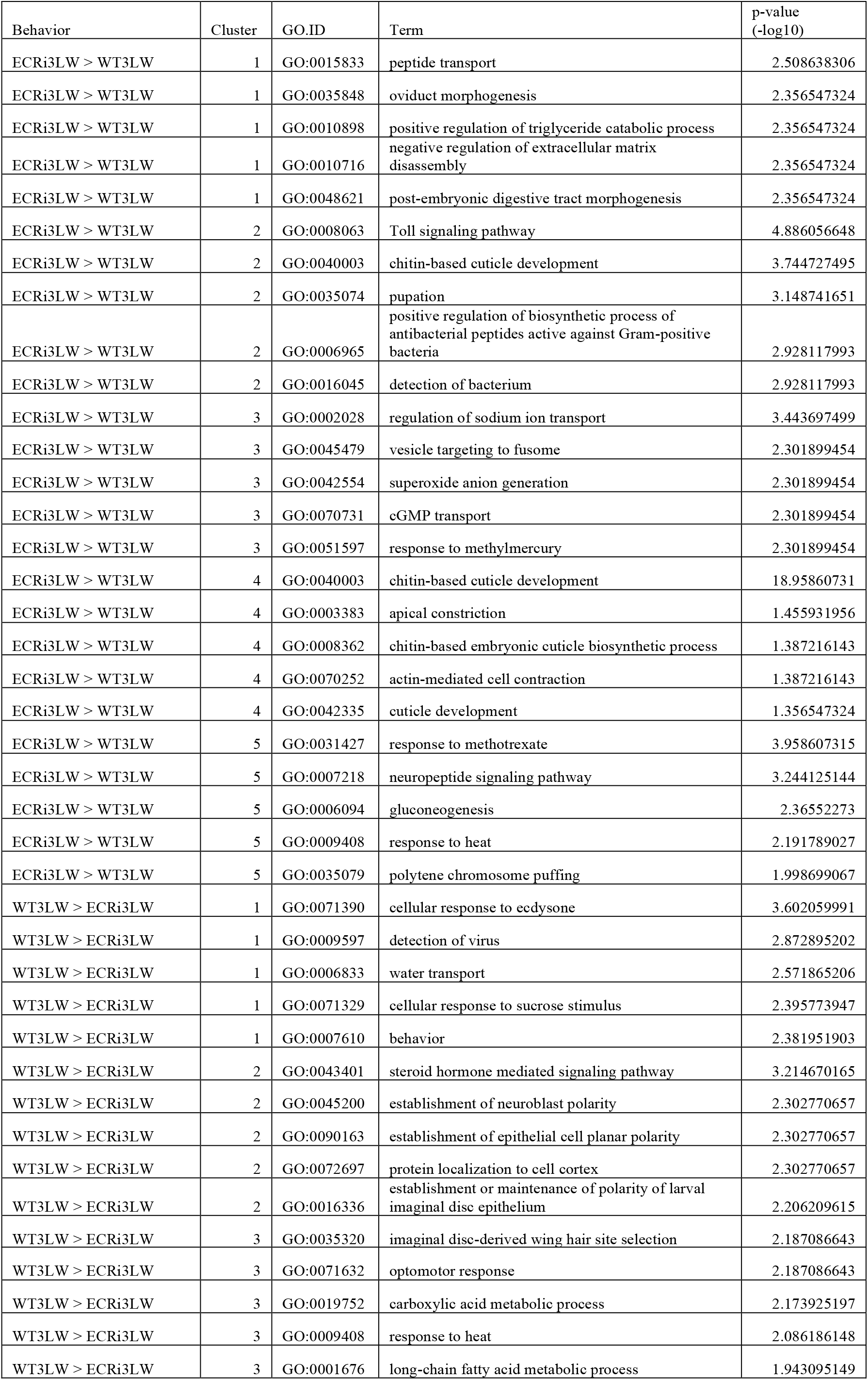
Gene Ontoogy Terms for EcR Clusters at −6hAPF (top five)

**Table S2:**
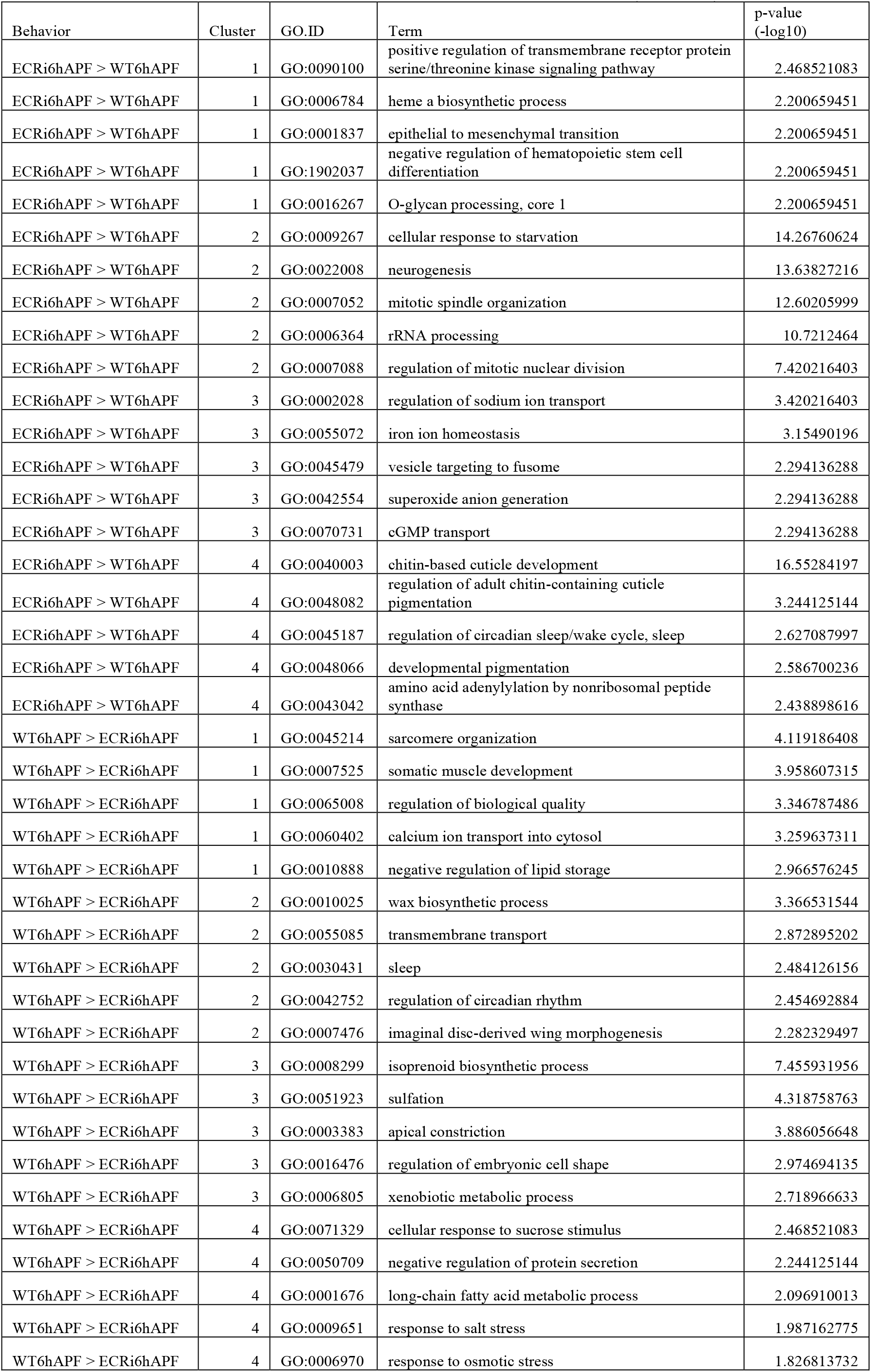
Gene Ontology Terms for EcR Clusters at +6hAPF (top five)

| Behavior | Cluster | GO.ID | Term | p-value<br>(-log10) |
| --- | --- | --- | --- | --- |
| ECRi6hAPF > WT6hAPF | 1 | GO:0090100 | positive regulation of transmembrane receptor protein serine/threonine kinase signaling pathway | 2.468521083 |
| ECRi6hAPF > WT6hAPF | 1 | GO:0006784 | heme a biosynthetic process | 2.200659451 |
| ECRi6hAPF > WT6hAPF | 1 | GO:0001837 | epithelial to mesenchymal transition | 2.200659451 |
| ECRi6hAPF > WT6hAPF | 1 | GO:1902037 | negative regulation of hematopoietic stem cell differentiation | 2.200659451 |
| ECRi6hAPF > WT6hAPF | 1 | GO:0016267 | O-glycan processing, core 1 | 2.200659451 |
| ECRi6hAPF > WT6hAPF | 2 | GO:0009267 | cellular response to starvation | 14.26760624 |
| ECRi6hAPF > WT6hAPF | 2 | GO:0022008 | neurogenesis | 13.63827216 |
| ECRi6hAPF > WT6hAPF | 2 | GO:0007052 | mitotic spindle organization | 12.60205999 |
| ECRi6hAPF > WT6hAPF | 2 | GO:0006364 | rRNA processing | 10.7212464 |
| ECRi6hAPF > WT6hAPF | 2 | GO:0007088 | regulation of mitotic nuclear division | 7.420216403 |
| ECRi6hAPF > WT6hAPF | 3 | GO:0002028 | regulation of sodium ion transport | 3.420216403 |
| ECRi6hAPF > WT6hAPF | 3 | GO:0055072 | iron ion homeostasis | 3.15490196 |
| ECRi6hAPF > WT6hAPF | 3 | GO:0045479 | vesicle targeting to fusome | 2.294136288 |
| ECRi6hAPF > WT6hAPF | 3 | GO:0042554 | superoxide anion generation | 2.294136288 |
| ECRi6hAPF > WT6hAPF | 3 | GO:0070731 | cGMP transport | 2.294136288 |
| ECRi6hAPF > WT6hAPF | 4 | GO:0040003 | chitin-based cuticle development | 16.55284197 |
| ECRi6hAPF > WT6hAPF | 4 | GO:0048082 | regulation of adult chitin-containing cuticle pigmentation | 3.244125144 |
| ECRi6hAPF > WT6hAPF | 4 | GO:0045187 | regulation of circadian sleep/wake cycle, sleep | 2.627087997 |
| ECRi6hAPF > WT6hAPF | 4 | GO:0048066 | developmental pigmentation | 2.586700236 |
| ECRi6hAPF > WT6hAPF | 4 | GO:0043042 | amino acid adenylation by nonribosomal peptide synthase | 2.438898616 |
| WT6hAPF > ECRi6hAPF | 1 | GO:0045214 | sarcomere organization | 4.119186408 |
| WT6hAPF > ECRi6hAPF | 1 | GO:0007525 | somatic muscle development | 3.958607315 |
| WT6hAPF > ECRi6hAPF | 1 | GO:0065008 | regulation of biological quality | 3.346787486 |
| WT6hAPF > ECRi6hAPF | 1 | GO:0060402 | calcium ion transport into cytosol | 3.259637311 |
| WT6hAPF > ECRi6hAPF | 1 | GO:0010888 | negative regulation of lipid storage | 2.966576245 |
| WT6hAPF > ECRi6hAPF | 2 | GO:0010025 | wax biosynthetic process | 3.366531544 |
| WT6hAPF > ECRi6hAPF | 2 | GO:0055085 | transmembrane transport | 2.872895202 |
| WT6hAPF > ECRi6hAPF | 2 | GO:0030431 | sleep | 2.484126156 |
| WT6hAPF > ECRi6hAPF | 2 | GO:0042752 | regulation of circadian rhythm | 2.454692884 |
| WT6hAPF > ECRi6hAPF | 2 | GO:0007476 | imaginal disc-derived wing morphogenesis | 2.282329497 |
| WT6hAPF > ECRi6hAPF | 3 | GO:0008299 | isoprenoid biosynthetic process | 7.455931956 |
| WT6hAPF > ECRi6hAPF | 3 | GO:0051923 | sulfation | 4.318758763 |
| WT6hAPF > ECRi6hAPF | 3 | GO:0003383 | apical constriction | 3.886056648 |
| WT6hAPF > ECRi6hAPF | 3 | GO:0016476 | regulation of embryonic cell shape | 2.974694135 |
| WT6hAPF > ECRi6hAPF | 3 | GO:0006805 | xenobiotic metabolic process | 2.718966633 |
| WT6hAPF > ECRi6hAPF | 4 | GO:0071329 | cellular response to sucrose stimulus | 2.468521083 |
| WT6hAPF > ECRi6hAPF | 4 | GO:0050709 | negative regulation of protein secretion | 2.244125144 |
| WT6hAPF > ECRi6hAPF | 4 | GO:0001676 | long-chain fatty acid metabolic process | 2.096910013 |
| WT6hAPF > ECRi6hAPF | 4 | GO:0009651 | response to salt stress | 1.987162775 |
| WT6hAPF > ECRi6hAPF | 4 | GO:0006970 | response to osmotic stress | 1.826813732 |

**Table S3:**
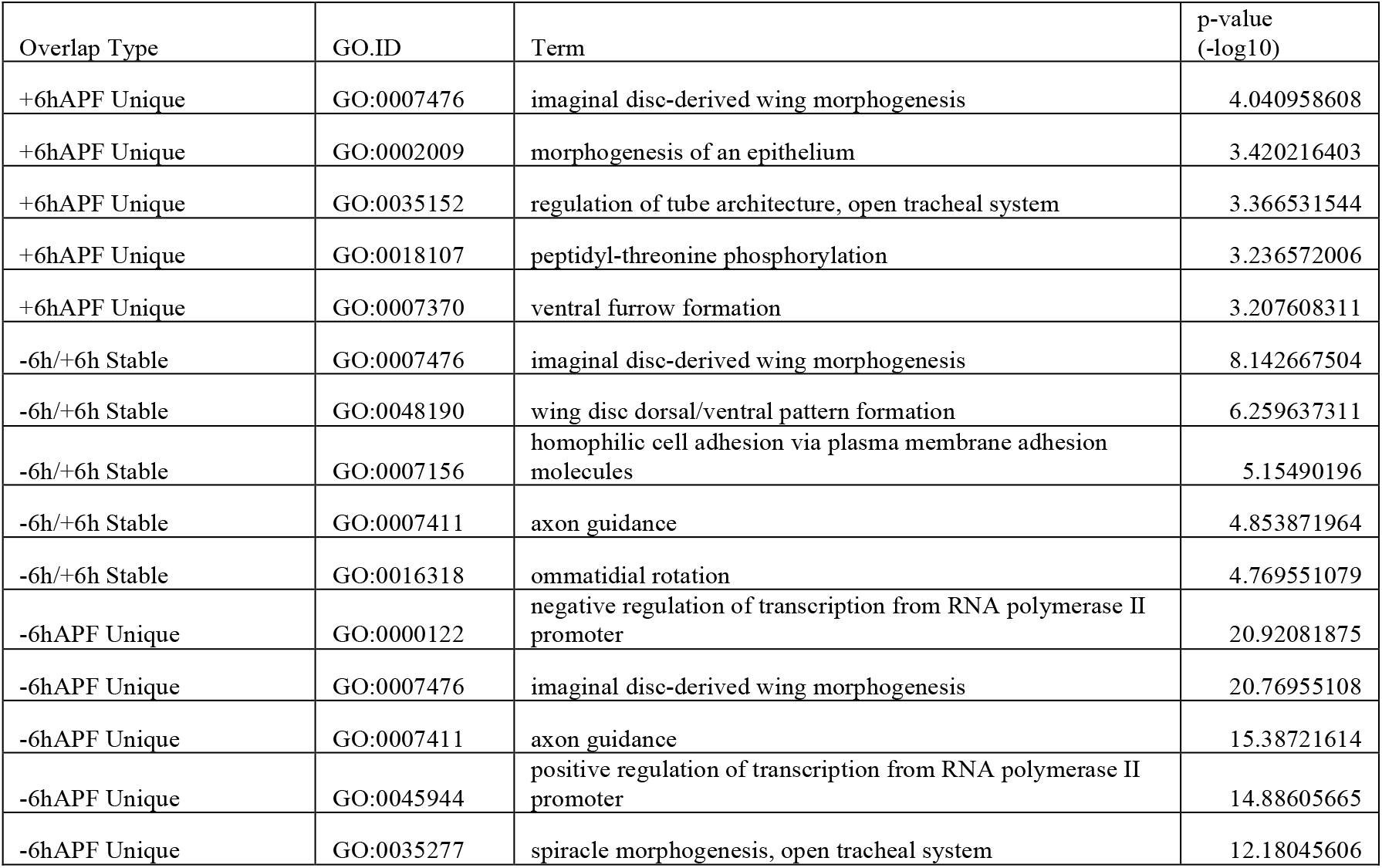
Gene Onto ogy Terms for EcR Binding Sites (top five)

| Overlap Type | GO.ID | Term | p-value<br>(-log10) |
| --- | --- | --- | --- |
| +6hAPF Unique | GO:0007476 | imaginal disc-derived wing morphogenesis | 4.040958608 |
| +6hAPF Unique | GO:0002009 | morphogenesis of an epithelium | 3.420216403 |
| +6hAPF Unique | GO:0035152 | regulation of tube architecture, open tracheal system | 3.366531544 |
| +6hAPF Unique | GO:0018107 | peptidyl-threonine phosphorylation | 3.236572006 |
| +6hAPF Unique | GO:0007370 | ventral furrow formation | 3.207608311 |
| -6h/+6h Stable | GO:0007476 | imaginal disc-derived wing morphogenesis | 8.142667504 |
| -6h/+6h Stable | GO:0048190 | wing disc dorsal/ventral pattern formation | 6.259637311 |
| -6h/+6h Stable | GO:0007156 | homophilic cell adhesion via plasma membrane adhesion molecules | 5.15490196 |
| -6h/+6h Stable | GO:0007411 | axon guidance | 4.853871964 |
| -6h/+6h Stable | GO:0016318 | ommatidial rotation | 4.769551079 |
| -6hAPF Unique | GO:0000122 | negative regulation of transcription from RNA polymerase II promoter | 20.92081875 |
| -6hAPF Unique | GO:0007476 | imaginal disc-derived wing morphogenesis | 20.76955108 |
| -6hAPF Unique | GO:0007411 | axon guidance | 15.38721614 |
| -6hAPF Unique | GO:0045944 | positive regulation of transcription from RNA polymerase II promoter | 14.88605665 |
| -6hAPF Unique | GO:0035277 | spiracle morphogenesis, open tracheal system | 12.18045606 |

